# The monothiol glutaredoxin Grx4 interacts with the Cryptococcus iron regulator Cirl and regulates iron homeostasis and virulence in the *Cryptococcus neoformans*

**DOI:** 10.1101/356774

**Authors:** Rodgoun Attarian, Guanggan Hu, Melissa Caza, Eddy Sanchez-Leon, Daniel Croll, Eunsoo Do, Horacio Bach, Tricia Missall, Jennifer Lodge, Won Hee Jung, James W. Kronstad

## Abstract

The acquisition of iron and the maintenance of iron homeostasis are important aspects of the virulence in the pathogenic fungus *Cryptococcus neoformans.* In this study, we identified the monothiol glutaredoxin Grx4 as a binding partner of Cir1, a master regulator of iron-responsive genes and virulence factor elaboration in *C. neoformans.* Monothiol glutaredoxins are important regulators of iron homeostasis because of their conserved roles in [2Fe-2S] cluster sensing and trafficking. We confirmed that Grx4 binds Cir1 and demonstrated that iron repletion promotes the relocalization of Grx4 from the nucleus to the cytoplasm. Nuclear retention is partially dependent on Cir1 and also influenced by treatment with the proteasome inhibitor bortezomib. Cir1 remains in the nucleus in both iron replete and iron limiting conditions. We also found that a *grx4*Δ mutant displayed iron-related phenotypes similar to those of a *cir1*Δ mutant, including poor growth upon iron deprivation. Importantly, a *grx4*Δ mutant was avirulent in mice, a phenotype consistent with observed defects in the key virulence determinants, capsule and melanin, and poor growth at 37°C. A comparative transcriptome analysis of a *grx4*Δ mutant and the WT strain in low iron and iron-replete conditions confirmed a central role for Grx4 in iron homeostasis. Dysregulation of iron-related metabolism was consistent with *grx4*Δ mutant phenotypes related to oxidative stress, mitochondrial function, and DNA repair. Overall, the phenotypes of the *grx4*Δ mutant and the RNA-Seq analysis support the hypothesis that Grx4 functions as a sensor of iron levels, in part through an interaction with Cir1, to extensively regulate iron homeostasis and contribute to virulence.

## Importance

Fungal pathogens cause life-threatening diseases in humans, particularly in immunocompromised people, and there is a tremendous need for a greater understanding of pathogenesis to support new therapies. One prominent fungal pathogen, *Cryptococcus neoformans*, causes menigitis in people suffering from HIV/AIDS. In the current study, we focused on characterizing mechanisms by which *C. neoformans* senses iron availability because iron is both a key nutrient and a signal for proliferation of the pathogen in vertebrate hosts. Specifically, we characterized a monothiol glutaredoxin protein Grx4 that functions as a sensor of iron, and that interacts with regulatory factors to control the ability of *C. neoformans* to cause disease. Grx4 regulates key virulence factors and a mutant is unable to cause disease in a mouse model of cryptococcosis. Overall, our study provides new insights into nutrient sensing and the role of iron in the pathogenesis of fungal diseases.

## Introduction

*Cryptococcus neoformans* is an opportunistic pathogen that causes life-threatening meningoencephalitis in immunocompromised people including those with HIV/AIDS (1–3). Despite the use of highly active antiretroviral therapy (HAART), there are still ~200,000 cases of cryptococcal meningoencephalitis per year, and the fungus is responsible for 15% of all AIDS-related deaths (4). This burden of disease underlines the urgent need to understand the mechanisms of fungus proliferation in vertebrate hosts as a foundation for identifying new drug and vaccine targets.

As with other pathogenic microbes, iron sensing and acquisition are important aspects of virulence for *C. neoformans* (5–8). Iron is important for *C. neoformans* both as a nutrient and as a signal to regulate the expression of the main virulence factor of the fungus, the polysaccharide capsule (5, 9). The abilities of *C. neoformans* to grow at the host body temperature of 37°C and to deposit melanin in the cell wall are also crucial for virulence (2, 6). To cause disease, the fungus must overcome nutritional immunity in which verterbrate hosts withhold iron to suppress pathogen growth (5, 11). *C. neoformans* employs various iron regulators and uptake mechanisms that contribute to virulence. These include heme uptake pathways as well as high and low affinity iron uptake systems (5, 12, 13). The use of heme as an iron source depends on an exported mannoprotein Cig1 and a cell surface reductase Fre2 (13, 14). High affinity uptake involves reduction of ferric iron (Fe^3+^) to the ferrous form (Fe^2+^) by cell surface reductases, with subsequent transport by a permease (Cft1) and ferroxidase (Cfo1) complex in the plasma membrane (5, 12, 14). The expression of these and other iron-related functions in *C. neoformans* is controlled by a GATA-type transcription factor Cir1 (cryptococcal iron regulator 1) and additional transcription factors including HapX (15, 16). Cir1 also integrates iron sensing and the regulation of iron uptake functions with the elaboration of virulence factors in *C. neoformans* (15).

Other fungi also use GATA-type transcriptional repressors with similar to Cir1 to regulate the expression of iron-responsive genes including *Schizosaccharomyces pombe* (Fep1), *Aspergillus sp.* (SreA), *Neurospora crassa* (SRE), and *Ustilago maydis* (Urbs1) (17–21). These transcription factors are characterized by one or two zinc finger motifs for DNA binding, and these flank a region containing four conserved cysteine residues. In contrast, the regulators of iron homeostasis in *Saccharomyces cerevisiae*, Aft1 and Aft2, are transcriptional activators (22, 23).

The mechanisms by which iron-responsive transcription factors in fungi sense intracellular iron levels and regulate iron homeostasis are best understood in *S. cerevisiae* and *S. pombe* (23–26). In these fungi, the transcription factors interact with monothiol glutaredoxins (GRXs) that participate in iron sensing and regulation (19, 24, 28–33). Monothiol GRXs are glutathione (GSH)-dependent proteins with a cysteine-glycine-phenylalanine-serine (CGFS) motif at the active site. These proteins are found in both prokaryotes and eukaryotes, and have emerged as key players in cellular redox and iron homeostasis (24, 35, 36). Recent studies in *S. cerevisiae* and *S. pombe* demonstrated essential roles for CGFS GRXs in intracellular iron homeostasis, iron trafficking and the maturation of [2Fe-2S] cluster proteins, and have established the proteins as novel [2Fe-2S] cluster-binding regulatory partners for transcription factors including Fep1 and Aft1 (24, 25, 27–33).

In this study, we demonstrate that the monothiol glutaredoxin Grx4 of *C. neoformans* interacts with Cir1. We also present evidence that Grx4 is involved in virulence and the maintenance of iron homeostasis. In particular, mutants lacking *GRX4* are defective for growth at the host temperature of 37°C and upon iron limitation. Along with defects in other virulence factors such as capsule and melanin, these findings account for the loss of virulence for the *grx4*Δ mutant in a murine model of cryptococcosis. Our results from transcriptional profiling by RNA-Seq further support a role for Grx4 in iron homeostasis through the regulation of functions for Fe-S cluster binding, heme biosynthesis, mitochondrial activities, and iron binding and uptake. These data support the conclusion that Grx4 is an important contributor to iron sensing and virulence in *C. neoformans.*

## Results

### The monothiol glutaredoxin Grx4 interacts with Cirl, a regulator of functions for iron uptake and virulence

Given that monothiol glutaredoxins (GRXs) make critical contributions to iron homeostasis in other fungi (24–27), we examined the genome sequence of the serotype A strain H99 of *C. neoformans* to identify candidate GRX proteins that could potentially interact with the key iron regulator Cir1. Specifically, we searched for orthologs of Grx3 and Grx4 from *S. cerevisiae* and Grx4 from *S. pombe* because these proteins are involved in iron regulation and homeostasis. A Blastp analysis identified a putative monothiol glutaredoxin encoded by the gene CNAG_02950 in *C. neoformans* that shared 63% amino acid sequence identity in the C-terminal region with Grx4 from *S. pombe*, 61% and 61% identity with Grx3 and Grx4 from *S. cerevisiae*, respectively, and 60% identity with Grx3 from *Homo sapiens* (Fig. Sl, Table Sl.). The C-terminal region of Grx4 from *C. neoformans* and the other monothiol glutaredoxin proteins contains a conserved glutaredoxin (GRX) domain with a signature “CGFS” (cysteine glycine phenylalanine serine) motif in the predicted active site (Fig. S1). Of note, the monothiol Grx domain with the CGFS active site motif is highly conserved and known to be required for [2Fe-2S] cluster binding and trafficking, and the regulation of iron homeostasis in fungi (24–34). The N-terminal region of Grx4 from *C. neoformans* also contained the WAXXC motif of the thioredoxin (TRX) domain found in monothiol glutaredoxins (24). Overall, the sequence analysis supports a possible role for Grx4 in iron-related processes in *C. neoformans.*

Grx3 and Grx4 in *S. cerevisiae* and Grx4 in *S. pombe* are known to interact with and to influence the activity of transcription factors that regulate iron homeostasis (24–34). We therefore employed the yeast two-hybrid assay to test the interaction between Grx4 and Cir1. As described in the Materials and Methods, the gene encoding Grx4 was synthesized and fused to the coding region for the Gal4 DNA binding domain (DBD) and the cDNA for Cir1 was fused with the Gal4 activating domain (AD) Yeast cells that harbored both DBD-Grx4 and AD-Cir1 were selected based on growth in the absence of leucine and tryptophan. The ability of these transformants to grow without uracil or histidine, and the expression of β-galactosidase, indicated a positive interaction between Grx4 and Cir1 (Fig. 1). The previously established interaction of Snf7 and Rim20 (37) was included as a positive control (Fig. 1). Overall, the results support the conclusion that Grx4 and Cirl interact.

**Figure 1.**
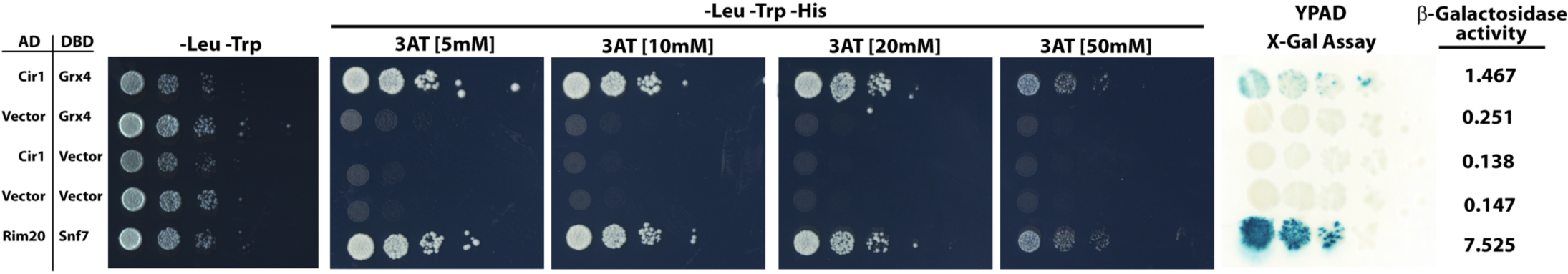
Grx4 interacts with Cirl. A yeast two-hybrid assay was used to examine the interaction of Grx4 and Cirl, DBD and AD indicate the Gal4 DNA binding and activation domains fused to Grx4 and Cirl, respectively. The vector designation indicates the empty vector control. All combinations of transformants grew in the absence of leucine (Leu) and tryptophan (Trp), confirming plasmid retention in the strains. Only yeast cells transformed with plasmids containing GRX4 and CIR1 grew in the absence of histidine (His), confirming an interaction to allow expression of HIS3. 3-Amino-1,2,4-trazole (3AT) was included at different concentrations to enhance the stringency of the *HIS3* selection. Qualitative and quantitative analyses of P-galactosidase activity were performed using X-gal or ONPG as a substrate, respectively and the quantitative numbers represent the mean values of three assays with the standard error of the mean provided in parentheses.

Grx4 co-localizes with Cirl in the nucleus upon iron limitation, but partially relocates to the cytoplasm upon iron repletion.

### Grx4 co-localizes with Cir1 in the nucleus upon iron limitation, but partially relocates to the cytoplasm upon iron repletion

The ability of Grx4 and Cir1 to interact suggested that the proteins would co-localize in the nucleus, and we therefore examined protein localization by tagging Grx4 with mCherry and Cir1 with GFP. Phenotypic assays indicated that the strains harboring the Cir1-GFP and Grx4-mCherry fusions behaved like the wild-type (WT) strain (Fig. S2), and the Grx4-mCherry and Cir1-GFP fusion proteins were detected as single bands by immunoblot analysis (Fig. S3). We cultured the strains carrying Grx4-mCherry and/or Cir1-GFP in low iron or iron-replete media for 5h at 30^°^C and examined fluorescence. As shown in Fig. 2A, both proteins were found in the nucleus in the low iron condition, and addition of iron as FeCl_3_ or heme resulted in partial relocalization of Grx4 to the cytoplasm (Fig. 2A). In contrast, Cir1 remained in the nucleus regardless of iron availability, and we confirmed the nuclear localization of Cir1-GFP by DAPI staining (Fig. 2A, Fig. S4). Quantitation of the relocation of Grx4-mCherry to the cytoplasm in response to iron and heme is shown in Figure 2B. We observed that the ratio of the nuclear to cytoplasmic signals from Grx4-mCherry shifted from ~4 fold to ~2 fold upon iron/heme repletion. We also examined the dependence of Grx4-mCherry localization on Cir1 by expressing the protein in a *cir1* deletion mutant (Fig. 2C). In this situation, Grx4 was observed to be mainly in the cytoplasm suggesting that the nuclear localization of Grx4 upon iron limitation is at least partially dependent on Cir1. An immunoblot analysis revealed that there was minimal decrease of protein levels for Grx4-mCherry in either the WT strain or *cir1* mutant after culturing the cells in YNB-BPS (low iron) or YNB-BPS+ FeCl_3_ for 5h (Fig. S3). As mentioned, Grx4-mCherry was distributed in the nucleus and cytoplasm in both WT and *cir1* mutant cells in the iron-replete condition (Fig. 2C). Interestingly, inclusion of the proteasome inhibitor bortezomib (BTZ) in the high iron condition appeared to enhance the level of the Grx4-mCherry protein that remained in the nucleus in both the WT strain and the *cir1* mutant (Fig. 2C). Although additional analyses are needed to investigate this result, it is possible that a proteasome-sensitive factor participates in the proper localization of Grx4 along with Cir1. BTZ treatment did not influence the localization of the Cir1-GFP protein in the WT strain regardless of iron availability (data not shown). Overall, the localization experiments further support an interaction between Grx4 and Cir1, and revealed that Grx4 localization is influenced by iron availability, Cir1, and the proteasome inhibitor BTZ.

**Figure 2.**
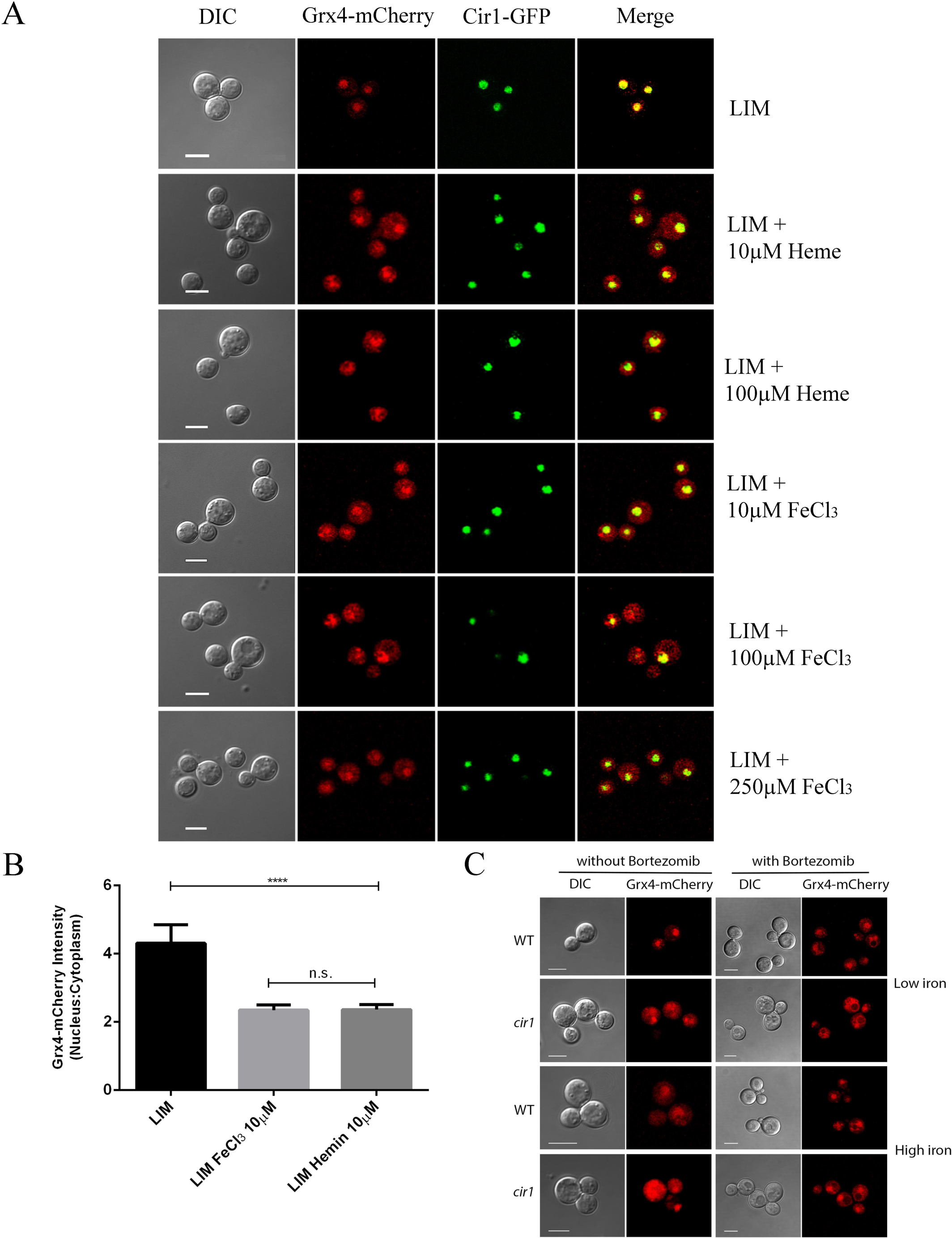
Grx4 is localized in nuclei in response to the iron limitation. A) Grx4-mCherry and Cirl-GFP were co-localized in nuclei in low iron conditions, but Grx4-mCherry shifted to cytosols with addition of iron supplement (FeCl_3_ or heme), while Cirl-GFP appears to stay in nuclei. Bar = 5 μM. B) Quantitation of the Grx4-mCherry signal in the nucleus and cytoplasm in response to different levels of iron, as described in the Materials and Methods. C) Deletion of *CIR1* caused mislocalization of Grx4-mCherry to the cytosol in the low iron condition. Grx4-mCherry was partially retained in nuclei in the high iron condition upon treatment with bortezomib, a proteasome inhibitor. Bar = 5 μM

#### Loss of Grx4 results in defects in the formation of major virulence factors and blocks cryptococcosis in mice

We next investigated the function of Grx4 in *C. neoformans* by constructing deletion mutants lacking the GRX domain of the *GRX4* gene. Interestingly, we were unable to delete the entire open reading frame, perhaps due to an essential function for the N-terminal TRX domain or an impact on an adjacent gene, but we did obtain two independent mutants lacking the C-terminal GRX domain (designated *grx4Δ-JL* and *grx4Δ-JK).* We also complemented the *grx4-JL* mutation with *GRX4.* The strains were initially compared with a mutant lacking Cir1 for phenotypes related to virulence. Specifically, we found that the *grx4Δ-JL* and *grx4Δ-JK* mutants displayed poor growth at 37^°^C, a phenotype shared with the *cir1* mutant (Fig. 3A) (15). The *grx4* mutants also had reduced production of melanin on medium containing L-DOPA as a substrate, and this was in contrast to the melanin production observed for the *cir1* mutant (Fig. 3B) (15). The polysaccharide capsule is a major virulence trait for *C. neoformans* and the loss of Grx4 also resulted in reduced capsule size, as does loss of Cir1 (15) (Fig. 3C, D). Taken together, these findings indicate that Grx4 is an important regulator of virulence factor production.

**Figure 3.**
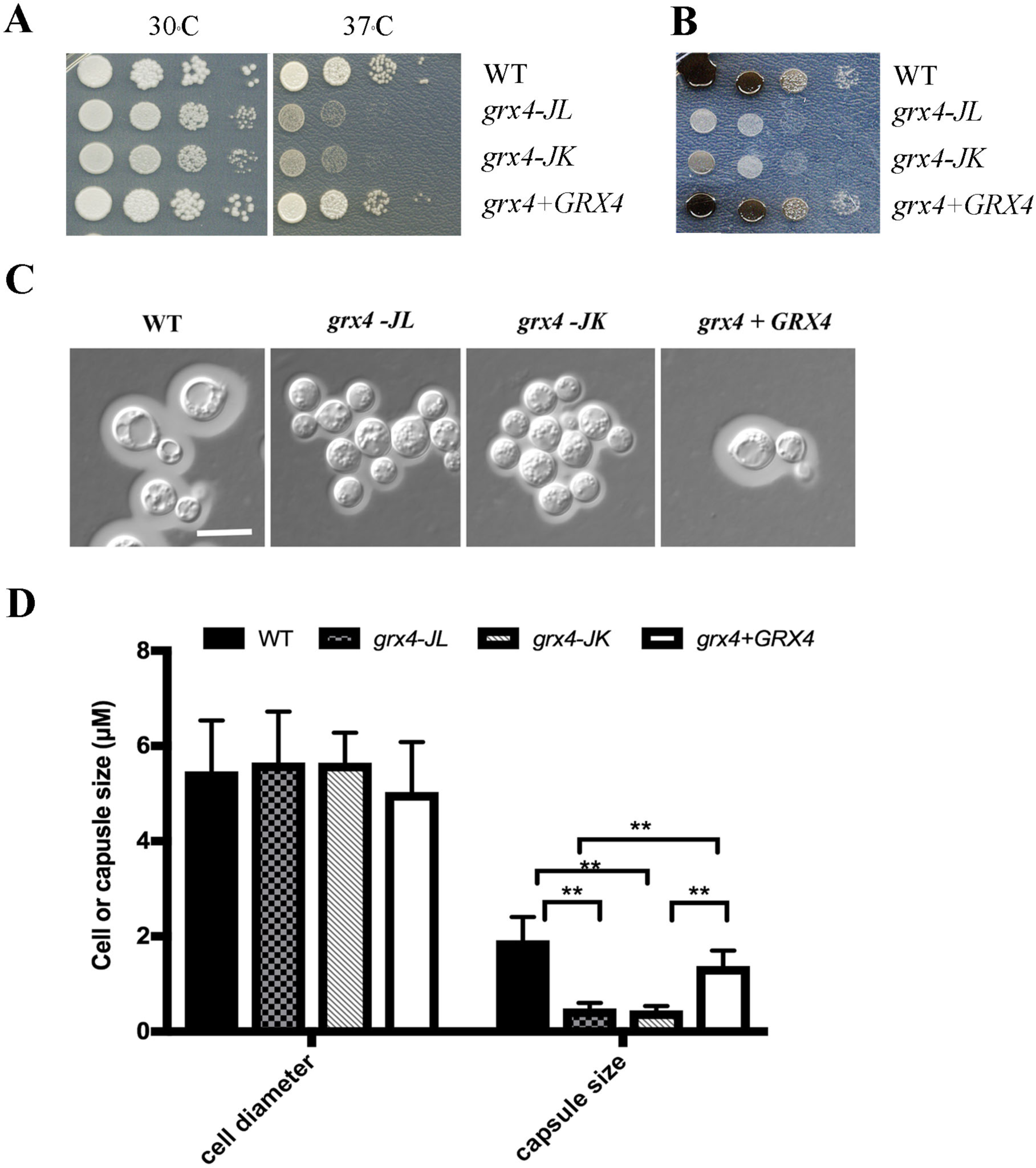
Disruption of *Grx4* influences the elaboration of three major virulence factors. A) The sensitivity of WT, two independent *grx4* mutants (one from the laboratory of Dr. J. Lodge (*grx4A-JL*) and the other from the laboratory of Dr. J. Kronstad *(grx4A-JK))* and the *GRX4* reconstituted strains to temperatures (30°C and 37°C) was examined. Aliquots of serial 10-fold dilutions of each strain were spotted on YPD plates and incubated for 2 days before being photographed. B) To examine melanin production, of aliquots serial 10-fold dilutions of each strain were spotted on L-DOPA plates and incubated for 2 days at 30°C before being photographed. C) Cells were grown in defined low-iron medium at 30°C for 48 h, and capsule formation was assessed by India ink staining for the indicated strains. Bar = 10 p,m. D) Fifty cells of each strain from the assays in A were measured to determine the cell diameter and capsule radius. Each bar represents the average of the 50 measurements with standard deviations. Statistical significance relative to the WT capsule size is indicated by two asterisks [Student’s t-test (P < 0.01)].

Our analysis of the impact of Grx4 on virulence-related phenotypes predicted that *grx4* mutants would be unable to cause disease in mice. To test this idea, we inoculated mice intranasally with cells of the WT strain, the *grx4Δ-JL* mutant or the complemented strain. All mice infected with the WT and complemented cells succumbed to infection by day 24, while the mice infected with the *grx4Δ-JL* mutant did not show disease symptoms and survived for the duration of the experiment (60 days, Fig. 4A). A more detailed examination of fungal burden in the infected mice revealed that the *grx4Δ-JL* mutant failed to accumulate in the brain, lung, liver, spleen, kidney and blood (Fig. 4B). Therefore, we conclude that Grx4 is required for the proliferation and/or survival of *C. neoformans* in a vertebrate host.

**Figure 4.**
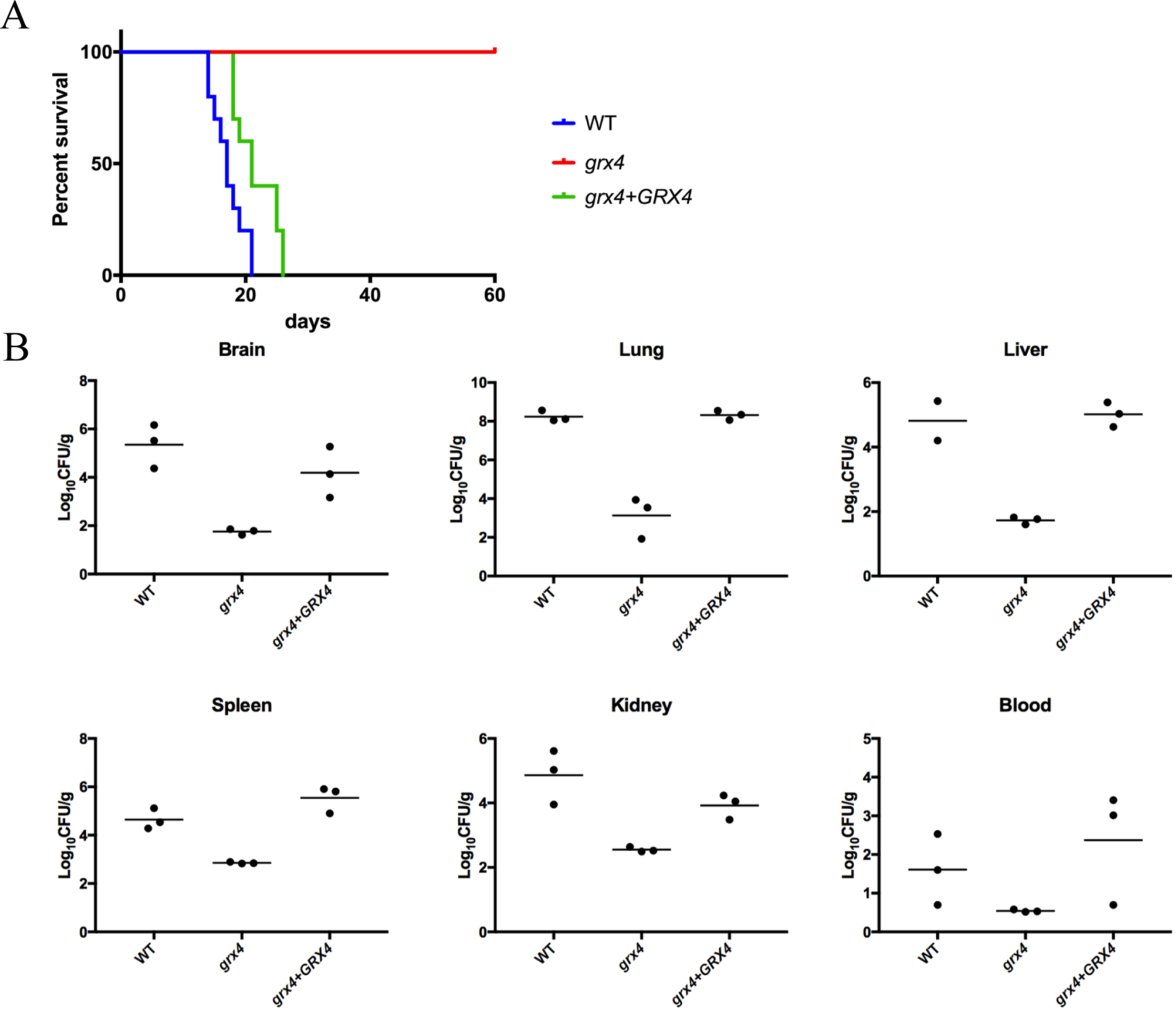
Grx4 is required for virulence in a mouse inhalation model. A) Ten female BALB/c mice were challenged by intranasal inoculation with 10^5^ cells of the WT strain (H99), the *grx4* mutant, or the *GRX4* reconstituted strain. The survival of the mice was monitored daily. Survival differences between groups of mice were evaluated by log rank tests. The P values for the mice infected with the WT and mutant strains were statistically significantly different (P <0.001). B) Distribution of fungal cells in the organs (brain, lung, live, spleen, kidney and blood) of mice infected by the inhalation method. Organs from mice infected with the WT, the grx4 mutant or the GRX4 reconstituted strain were collected at human endpint of the experiment, and fungal burdens were monitored in organs by determining colony-forming units (CFU) upon plating on YPD medium. Three mice for each strain were used for the experiments, and horizontal bars in each graph represent an average CFUs. In all organs, differences of fungal burdens between the *grx4* mutant and the WT strain, and between the *grx4* mutant and the reconstituted strain, were statistical significance (p < 0.05).

#### Grx4 is required for growth on low iron media

Given that Grx4 is critical for the virulence of *C. neoformans* and shares some phenotypes with Cir1, we hypothesized that Grx4 might also contribute along with Cir1 to iron homeostasis. We therefore examined the ability of the *grx4* mutants to proliferate on media with low and high concentrations of iron and heme (Fig. 5). We found that the mutants showed poor proliferation on solid media with FeCl_3_, FeSO_4_ or heme as iron sources. The impaired growth was particularly notable at low iron levels where the cells are dependent on high affinity iron uptake. Similar defects in proliferation for the mutants were observed for cultures in liquid media with reduced iron availability. We also examined the influence of iron on the *grx4* mutants in more detail by testing their sensitivity to iron chelators (e.g., curcumin and ferrozine), as well as their ability to proliferate in the presence of elevated iron concentrations (Fig. 6A-C). These experiments revealed that the mutants were sensitive to curcumin and high concentrations of ferrozine. Heme and FeEDTA partially rescued the inhibitory effects of curcumin or ferrozine, respectively (Fig. 6A, B). Elevated iron in the culture medium slightly impaired the proliferation of the mutants (Fig. 6C). Overall, these results indicate that Grx4 participates in iron homeostasis in *C. neoformans*, a role that is consistent with its interaction with Cir1.

**Figure 5.**
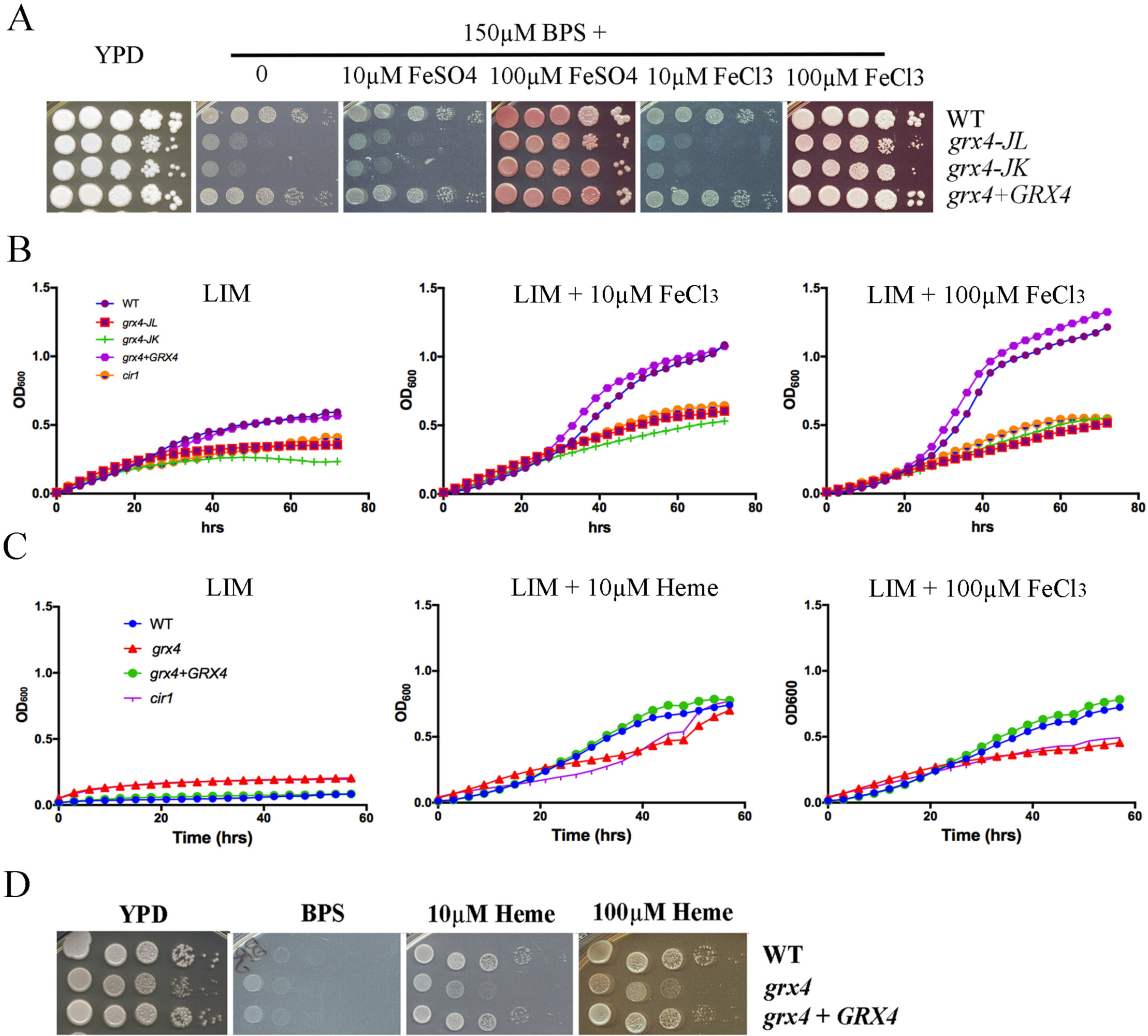
Grx4 is required for robust growth on inorganic iron or heme as the sole iron source. A) Tenfold serial dilutions of each strain (labeled on the right) were spotted on the indicated media and the plates were incubated at 30°C for 5 days before being photographed. B) Cells of the WT, the *grx4* mutant, and the *GRX4* reconstituted strain were inoculated into liquid YNB medium plus 150 μM BPS without and with supplementation with FeCl_3_ as the iron source. The cultures were incubated at 30°C, and OD_600_s were measured. The *cir1* mutant strain was included for comparison with the *grx4* strains. C) The indicated strains (the WT, the *grx4* mutant, and the *GRX4* reconstituted strain) were also tested for growth in the LIM (liquid YNB medium plus 150 μM BPS) without and with supplementation with heme by the same method used for panel B. D) Tenfold serial dilutions of each strain were spotted on the indicated media and the plates were incubated at 30°C for 2 days before being photographed

**Figure 6.**
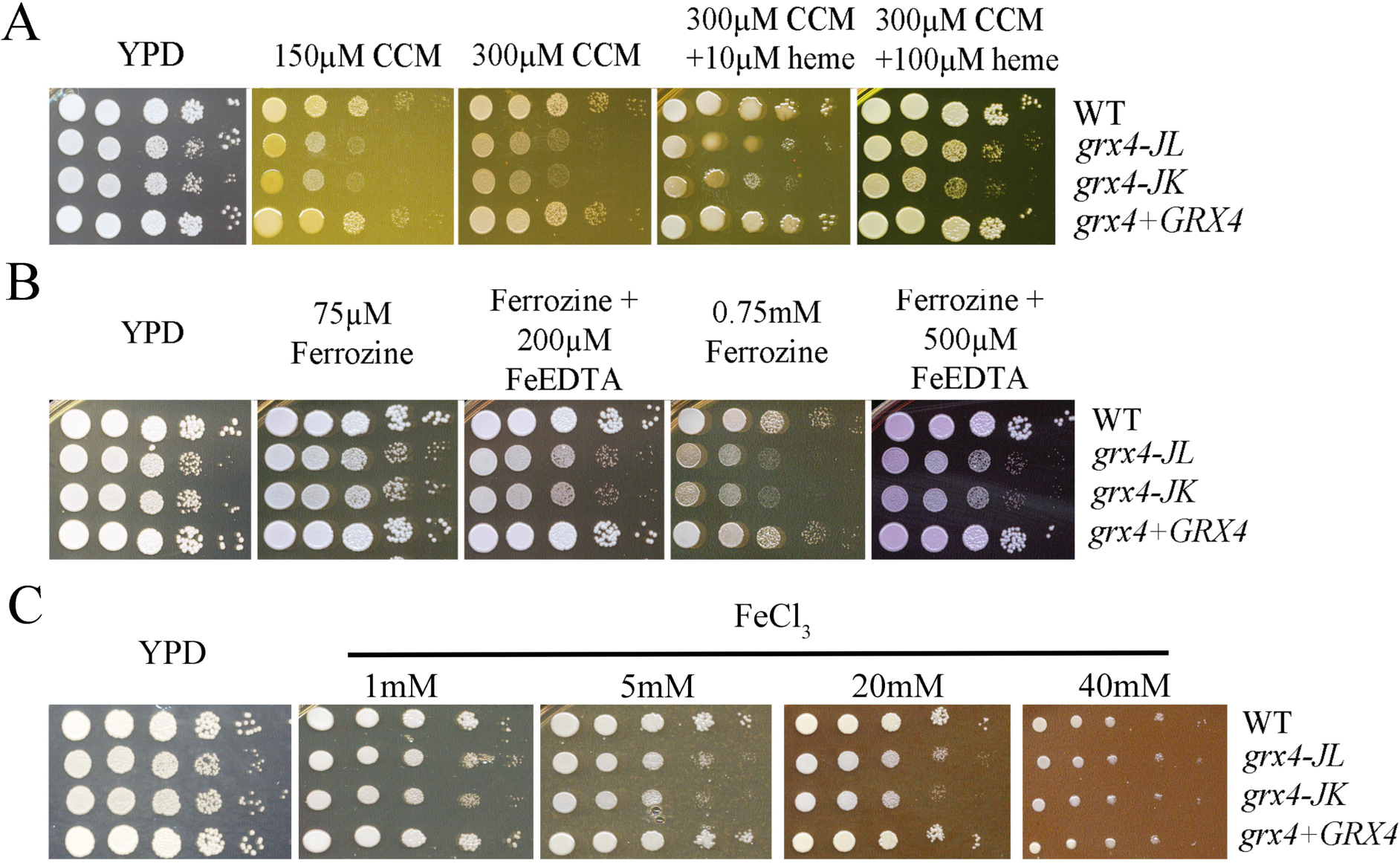
Grx4 is involved in iron homeostasis. A) and B) Disruption of *GRX4* leads to increased sensitivity to the iron - chelating drugs, curcumin and ferrozine. A) Tenfold serial dilutions of each strain, including the WT, two independent *grx4* mutants, and the *GRX4* complemented strains (without iron starvation), were spotted on the YPD plates with or without curcumin at a concentration of either 150 or 300 μM, supplemented with 0, 10, or 100 μM of heme as the iron source. The plates were incubated at 30 °C for 2 days before being photographed. B) Tenfold serial dilutions of each strain without iron starvation were spotted on the YPD plates with or without 75μM or 750μM ferrozine supplemented with 0, or indicated amount of FeEDTA (Jung et al., 2006). The plates were incubated at 30 °C for 2 days before being photographed. C) Disruption of *GRX4* caused increased sensitivity to iron overload. Tenfold serial dilutions of each strain grown in YPD medium overnight were spotted onto YPD supplemented with either 1mM, or 5mM, or 20mm, or 40mM FeCl_3_. The plates were incubated at 30 °C for 2 days before being photographed. D) and E). Increased iron accumulation in the *grx4* and *cir1* mutant strains. Total cellular iron levels in the WT, *grx4* and *cir1* mutant strains were determined using ICP analysis. D) The samples were collected from the cells of each strain grown under low iron condition (YNB + 150 μM BPS) at 30 °C overnight. E) The samples were collected from the cells of each strain grown under high iron condition (YNB + 150 μM BPS + 100 μM FeCl_3_) at 30 °C overnight. Values indicate weight per gram of dry cells.

#### Transcriptional profiling supports a role for Grx4 in the regulation of iron-dependent processes

The glutaredoxin Grx4 in *S. pombe* is known to co-regulate genes for iron acquisition with Fep1, an ortholog of Cir1, and to interact with Php4, a regulator of functions that use iron (27, 29, 30, 33, 34). We therefore performed an RNA-Seq analysis to assess the impact of loss of Grx4 on the transcriptomes of *C. neoformans* cells grown in low and high iron conditions (Fig. 7). As shown in Table 1, loss of Grx4 had an impact on transcript levels for several hundred genes in both low and high iron conditions. That is, ~647 genes were up-regulated in the *grx4* mutant compared with the WT strain in the low iron condition, and 736 genes were up-regulated in the high iron condition. We also found that 404 genes were down-regulated in the *grx4* mutant in the low iron condition versus 358 genes in high iron (Table 1). The lists of up-regulated or down-regulated transcripts in both conditions are presented in Tables S2 and S3, respectively.

**Table 1.** Number of genes differentially regulated by iron availability and/or Grx4

| Comparison | up-regulated |  |  |  | down-regulated |  |  |  |
| --- | --- | --- | --- | --- | --- | --- | --- | --- |
|  | 2-fold | 5-fold | 10-fold | total | 2-fold | 5-fold | 10-fold | total |
| WT (low iron vs high iron) | 525 | 16 | 0 | 541 | 328 | 53 | 27 | 408 |
| *grx4* vs WT, low iron | 498 | 99 | 50 | 647 | 317 | 62 | 25 | 404 |
| *grx4* vs WT, high iron | 580 | 100 | 56 | 736 | 290 | 52 | 16 | 358 |
| *grx4* (low iron vs high iron) | 100 | 2 | 0 | 102 | 237 | 11 | 0 | 248 |

**Figure 7.**
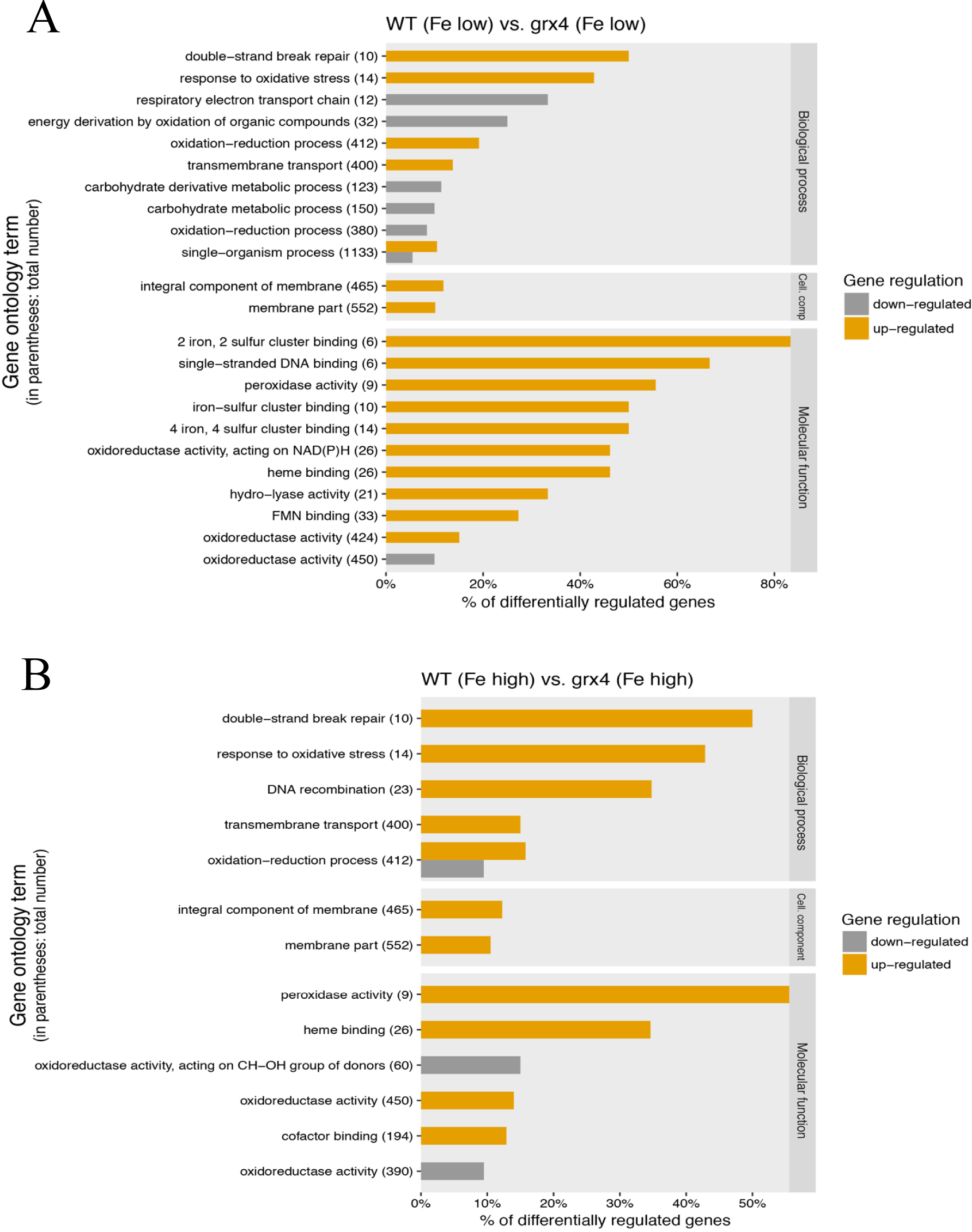
Impact of loss of Grx4 on the transcripts for specific gene ontology categories. A) and B) Gene ontology (GO) enrichment analysis of the differentially expressed genes between WT and *grx4* strains under low iron (A) and high iron (B) conditions (with total gene numbers within each functional category, %: percentage of genes showing differential expression).

An analysis of gene ontology terms for biological processes revealed that the highest percentages of differentially expressed and up-regulated genes (top 3) in the low-iron condition were in double-strand break repair, response to oxidative stress and oxidation-reduction processes (Fig. 7A). Genes for respiratory electron transport chain, energy derivation by oxidation of organic compounds and carbohydrate derivative metabolic process were the top categories for the down-regulated transcripts (Fig. 7A). For the iron-replete condition, the top categories for up-regulated transcripts were double-strand break repair, response to oxidative stress, and DNA recombination, while the single category noted for down-regulated transcripts was oxidation-reduction process (Fig. 7B). Notable GO terms for molecular function also implicated Grx4 in the regulation of 2Fe-2S cluster binding, single-stranded DNA binding, peroxidase activity, oxidoreductase activity, and heme binding (Fig. 7B). The influence of *C. neoformans* on DNA-related processes and iron-dependent mitochondrial functions related to respiration and oxidative phosphorylation was also highlighted by STRING analysis (38) (Fig. S5). As a whole, these results, along with the interaction of Grx4 with the iron regulator Cir1 (Figs. 1 and 2) and the influence of the *grx4* deletion on growth on low iron media (Fig. 5), highlight the important role that Grx4 plays in iron-related processes in *C. neoformans.* We manually examined the categories of genes whose transcript levels are influenced by Grx4 and compiled a table to illustrate selected functions including iron transport, heme biosynthesis, mitochondrial and cytosolic iron sulfur cluster assembly, iron sulfur containing proteins, electron transport, mitochondrial functions, DNA repair, ergosterol metabolism and oxidative stress (Table 2). We also performed qRT-PCR to compare the expression of genes encoding the functions related to iron *(CIR1, LAC1* and *FRE3)* in WT and *grx4* mutant cells, and found results consistent with the RNAseq data (Fig. S6). In general, these results highlighted the negative impact of Grx4 on the regulation of iron-dependent functions (Table 2), similar to the influence of the monothiol glutaredoxin Grx4 with the transcription factor Php4 in *S. pombe* (27, 33, 34).

**Table 2.**
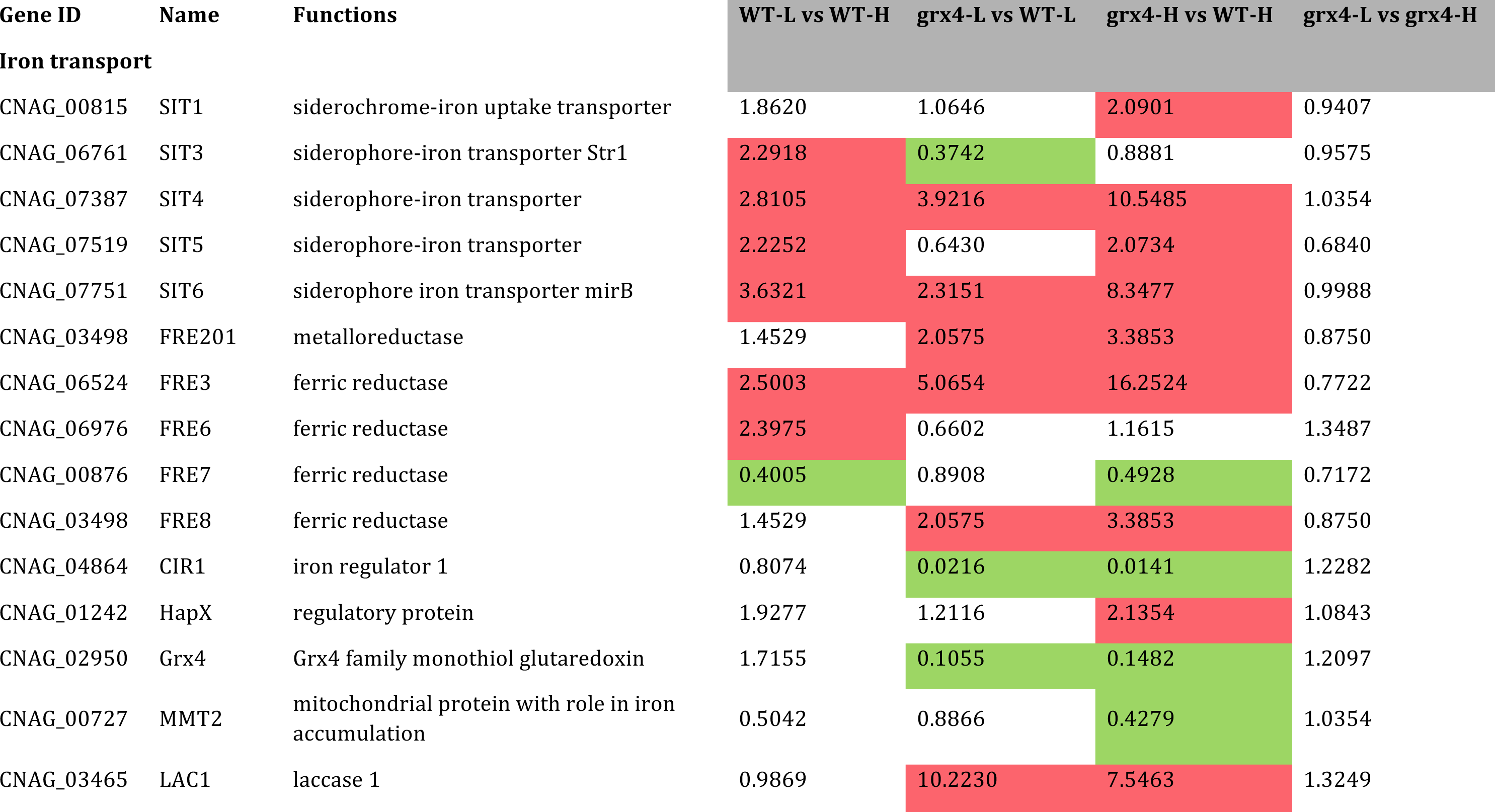
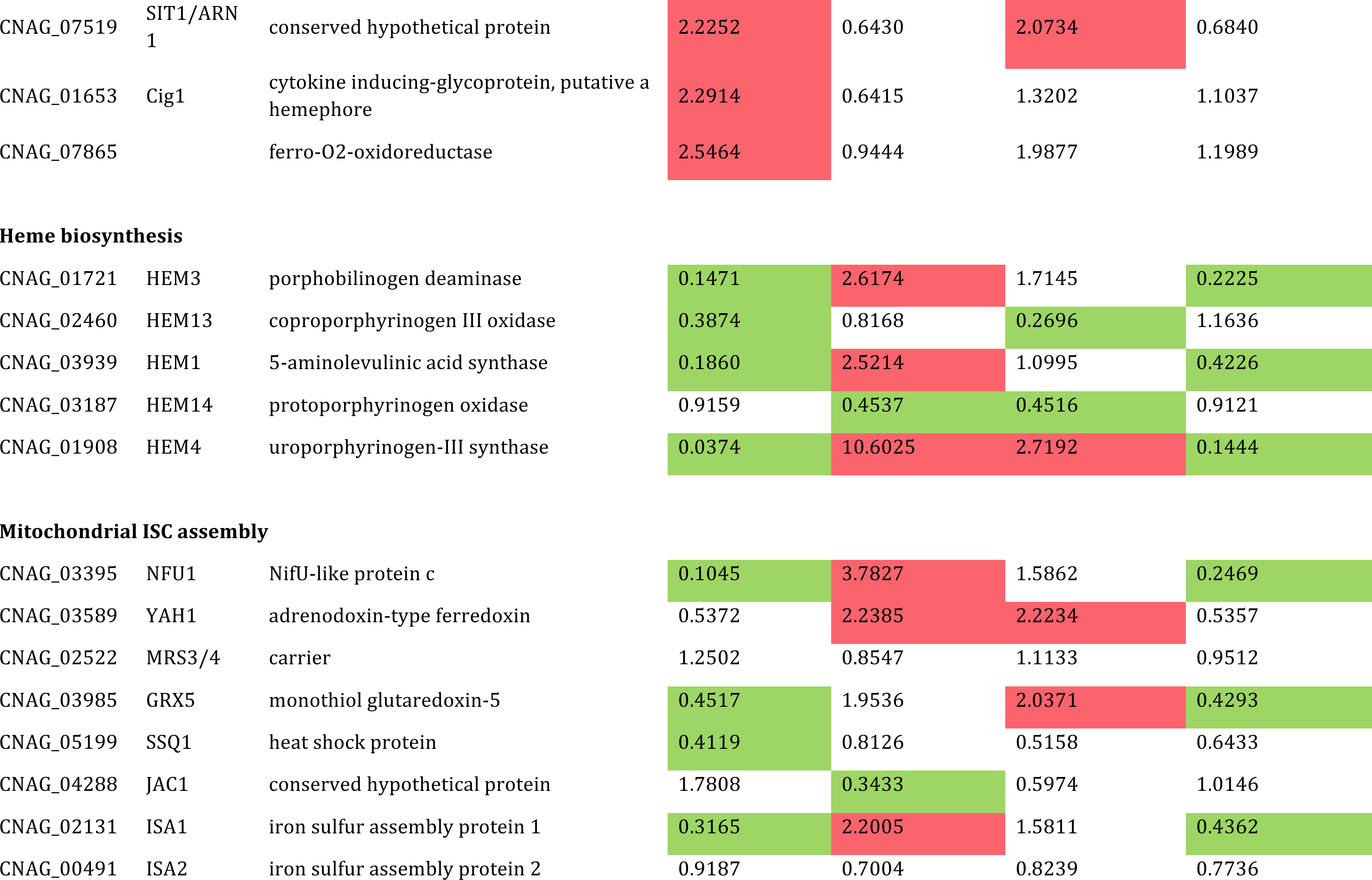
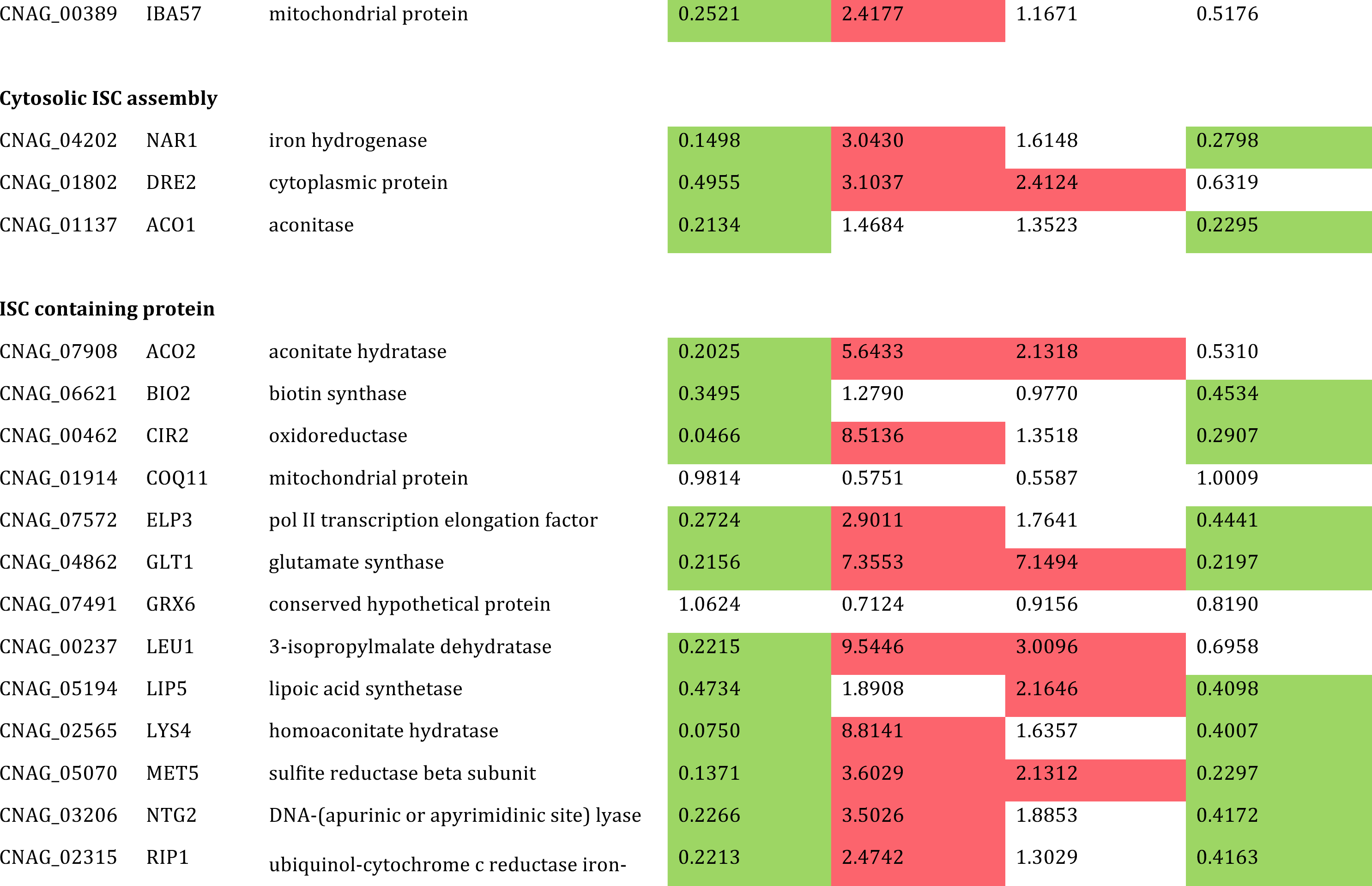
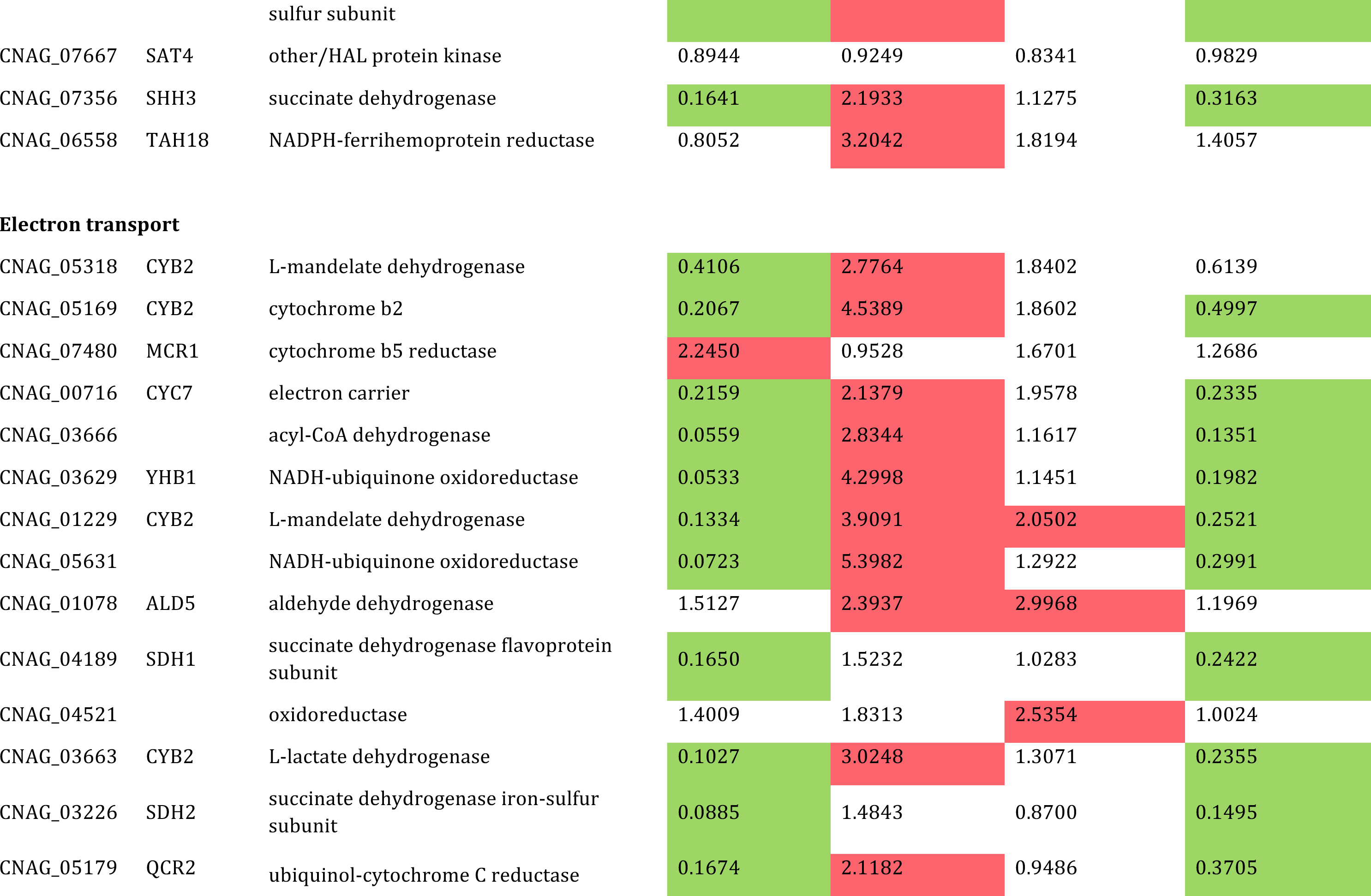
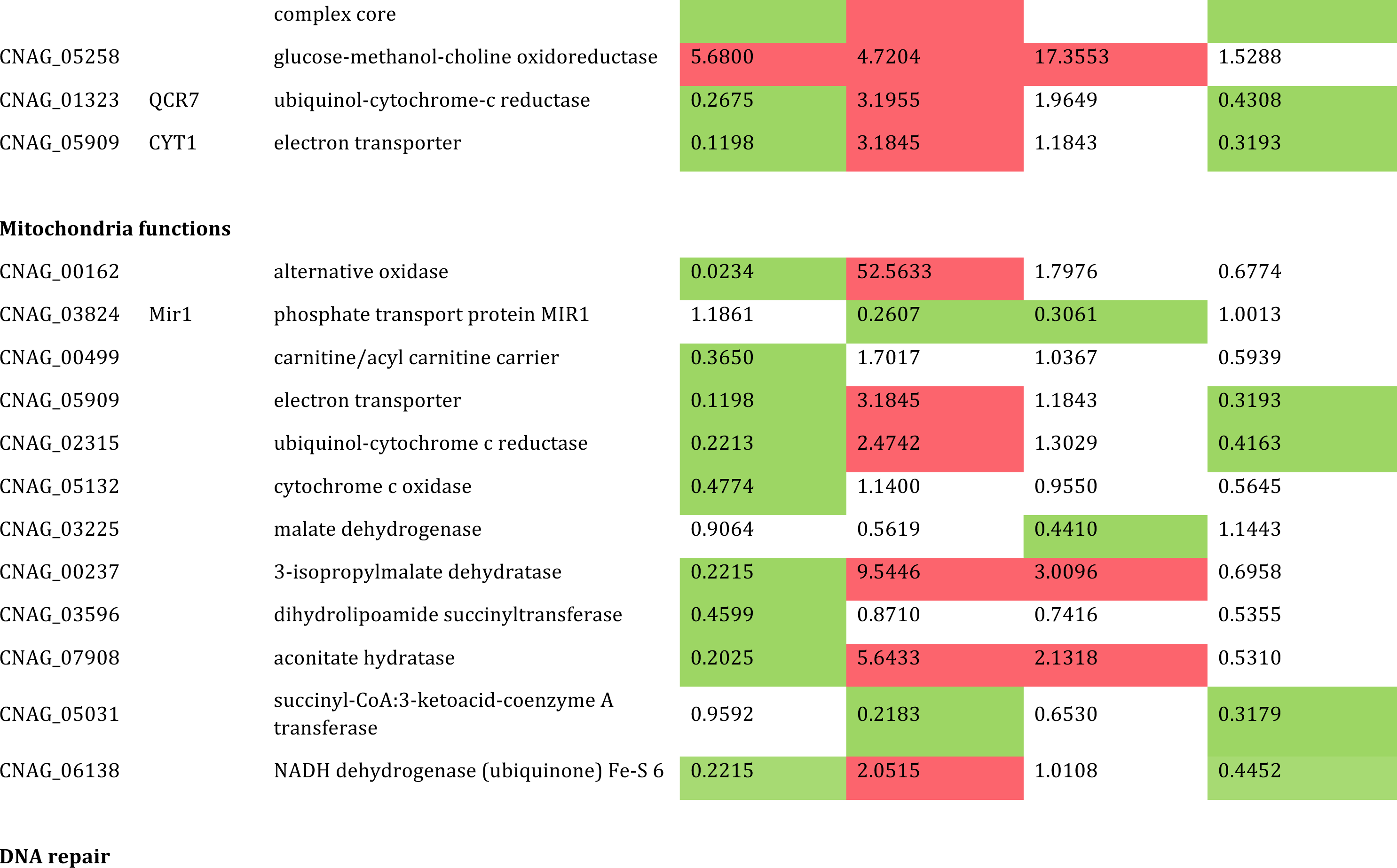
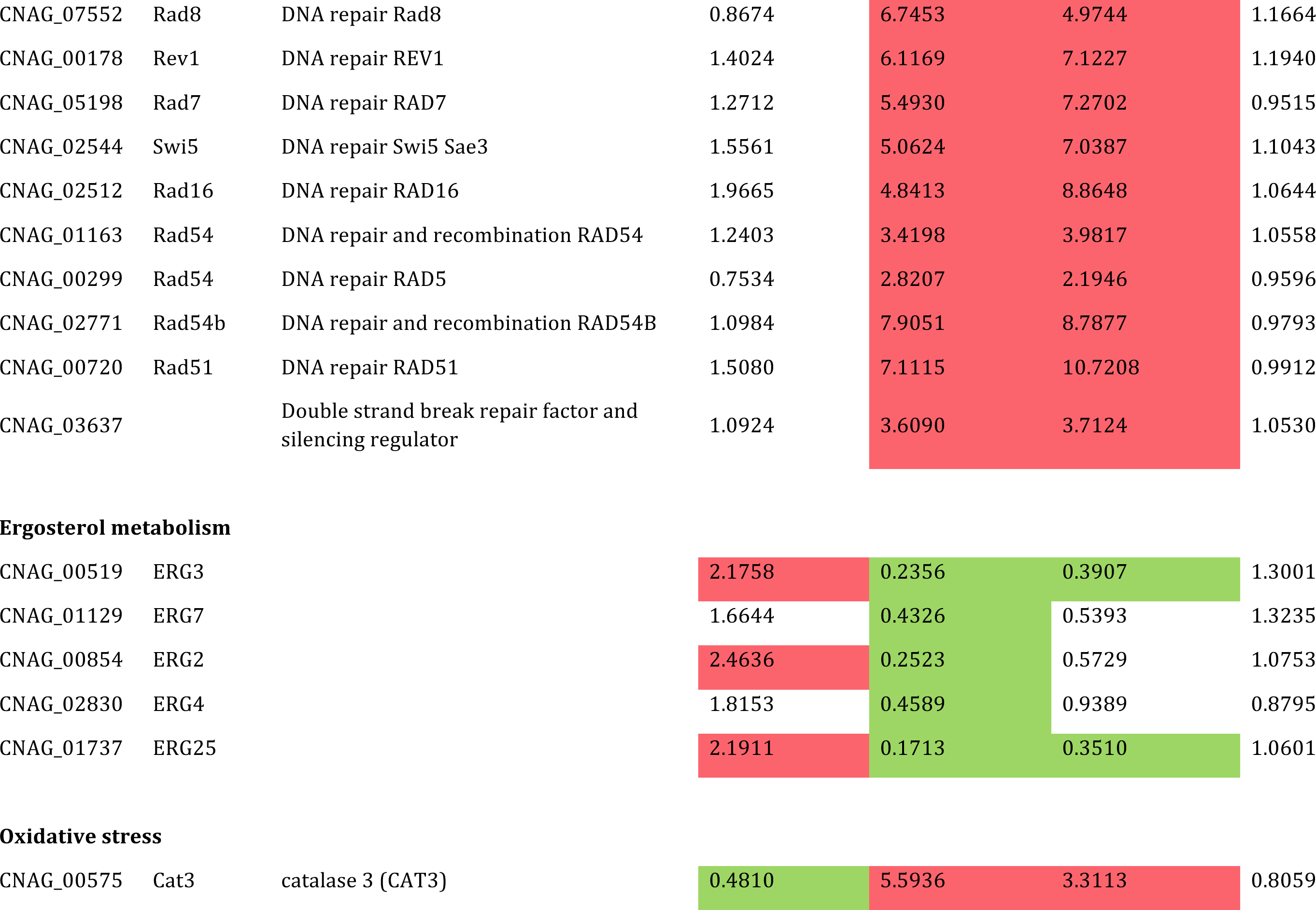
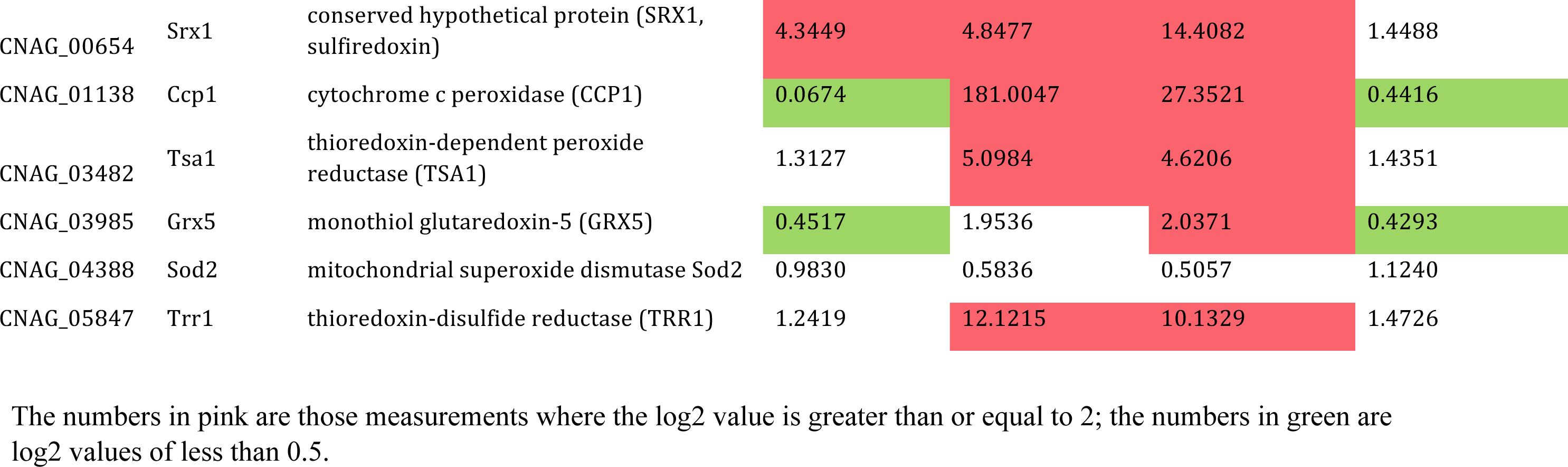
Regulation by Grx4 of genes encoding iron transport, iron homeostasis and mitochondrial functions

| Gene ID | Name | Functions | WT-L vs WT-H | grx4-L vs WT-L | grx4-H vs WT-H | grx4-L vs grx4-H |
| --- | --- | --- | --- | --- | --- | --- |
| **Iron transport** |  |  |  |  |  |  |
| CNAG_00815 | SIT1 | siderochrome-iron uptake transporter | 1.8620 | 1.0646 | 2.0901 | 0.9407 |
| CNAG_06761 | SIT3 | siderophore-iron transporter Str1 | 2.2918 | 0.3742 | 0.8881 | 0.9575 |
| CNAG_07387 | SIT4 | siderophore-iron transporter | 2.8105 | 3.9216 | 10.5485 | 1.0354 |
| CNAG_07519 | SIT5 | siderophore-iron transporter | 2.2252 | 0.6430 | 2.0734 | 0.6840 |
| CNAG_07751 | SIT6 | siderophore iron transporter mirB | 3.6321 | 2.3151 | 8.3477 | 0.9988 |
| CNAG_03498 | FRE201 | metalloreductase | 1.4529 | 2.0575 | 3.3853 | 0.8750 |
| CNAG_06524 | FRE3 | ferric reductase | 2.5003 | 5.0654 | 16.2524 | 0.7722 |
| CNAG_06976 | FRE6 | ferric reductase | 2.3975 | 0.6602 | 1.1615 | 1.3487 |
| CNAG_00876 | FRE7 | ferric reductase | 0.4005 | 0.8908 | 0.4928 | 0.7172 |
| CNAG_03498 | FRE8 | ferric reductase | 1.4529 | 2.0575 | 3.3853 | 0.8750 |
| CNAG_04864 | CIR1 | iron regulator 1 | 0.8074 | 0.0216 | 0.0141 | 1.2282 |
| CNAG_01242 | HapX | regulatory protein | 1.9277 | 1.2116 | 2.1354 | 1.0843 |
| CNAG_02950 | Grx4 | Grx4 family monothiol glutaredoxin | 1.7155 | 0.1055 | 0.1482 | 1.2097 |
| CNAG_00727 | MMT2 | mitochondrial protein with role in iron accumulation | 0.5042 | 0.8866 | 0.4279 | 1.0354 |
| CNAG_03465 | LAC1 | laccase 1 | 0.9869 | 10.2230 | 7.5463 | 1.3249 |
| CNAG_07519 | SIT1/ARN<br>1 | conserved hypothetical protein | 2.2252 | 0.6430 | 2.0734 | 0.6840 |
| CNAG_01653 | Cig1 | cytokine inducing-glycoprotein, putative a<br>hemephore | 2.2914 | 0.6415 | 1.3202 | 1.1037 |
| CNAG_07865 |  | ferro-O2-oxidoreductase | 2.5464 | 0.9444 | 1.9877 | 1.1989 |
| Heme biosynthesis |  |  |  |  |  |  |
| CNAG_01721 | HEM3 | porphobilinogen deaminase | 0.1471 | 2.6174 | 1.7145 | 0.2225 |
| CNAG_02460 | HEM13 | coproporphyrinogen III oxidase | 0.3874 | 0.8168 | 0.2696 | 1.1636 |
| CNAG_03939 | HEM1 | 5-aminolevulinic acid synthase | 0.1860 | 2.5214 | 1.0995 | 0.4226 |
| CNAG_03187 | HEM14 | protoporphyrinogen oxidase | 0.9159 | 0.4537 | 0.4516 | 0.9121 |
| CNAG_01908 | HEM4 | uroporphyrinogen-III synthase | 0.0374 | 10.6025 | 2.7192 | 0.1444 |
| Mitochondrial ISC assembly |  |  |  |  |  |  |
| CNAG_03395 | NFU1 | NifU-like protein c | 0.1045 | 3.7827 | 1.5862 | 0.2469 |
| CNAG_03589 | YAH1 | adrenodoxin-type ferredoxin | 0.5372 | 2.2385 | 2.2234 | 0.5357 |
| CNAG_02522 | MRS3/4 | carrier | 1.2502 | 0.8547 | 1.1133 | 0.9512 |
| CNAG_03985 | GRX5 | monothiol glutaredoxin-5 | 0.4517 | 1.9536 | 2.0371 | 0.4293 |
| CNAG_05199 | SSQ1 | heat shock protein | 0.4119 | 0.8126 | 0.5158 | 0.6433 |
| CNAG_04288 | JAC1 | conserved hypothetical protein | 1.7808 | 0.3433 | 0.5974 | 1.0146 |
| CNAG_02131 | ISA1 | iron sulfur assembly protein 1 | 0.3165 | 2.2005 | 1.5811 | 0.4362 |
| CNAG_00491 | ISA2 | iron sulfur assembly protein 2 | 0.9187 | 0.7004 | 0.8239 | 0.7736 |
| CNAG_00389 | IBA57 | mitochondrial protein | 0.2521 | 2.4177 | 1.1671 | 0.5176 |
| **Cytosolic ISC assembly** |  |  |  |  |  |  |
| CNAG_04202 | NAR1 | iron hydrogenase | 0.1498 | 3.0430 | 1.6148 | 0.2798 |
| CNAG_01802 | DRE2 | cytoplasmic protein | 0.4955 | 3.1037 | 2.4124 | 0.6319 |
| CNAG_01137 | ACO1 | aconitase | 0.2134 | 1.4684 | 1.3523 | 0.2295 |
| **ISC containing protein** |  |  |  |  |  |  |
| CNAG_07908 | ACO2 | aconitate hydratase | 0.2025 | 5.6433 | 2.1318 | 0.5310 |
| CNAG_06621 | BIO2 | biotin synthase | 0.3495 | 1.2790 | 0.9770 | 0.4534 |
| CNAG_00462 | CIR2 | oxidoreductase | 0.0466 | 8.5136 | 1.3518 | 0.2907 |
| CNAG_01914 | COQ11 | mitochondrial protein | 0.9814 | 0.5751 | 0.5587 | 1.0009 |
| CNAG_07572 | ELP3 | pol II transcription elongation factor | 0.2724 | 2.9011 | 1.7641 | 0.4441 |
| CNAG_04862 | GLT1 | glutamate synthase | 0.2156 | 7.3553 | 7.1494 | 0.2197 |
| CNAG_07491 | GRX6 | conserved hypothetical protein | 1.0624 | 0.7124 | 0.9156 | 0.8190 |
| CNAG_00237 | LEU1 | 3-isopropylmalate dehydratase | 0.2215 | 9.5446 | 3.0096 | 0.6958 |
| CNAG_05194 | LIP5 | lipoic acid synthetase | 0.4734 | 1.8908 | 2.1646 | 0.4098 |
| CNAG_02565 | LYS4 | homoaconitate hydratase | 0.0750 | 8.8141 | 1.6357 | 0.4007 |
| CNAG_05070 | MET5 | sulfite reductase beta subunit | 0.1371 | 3.6029 | 2.1312 | 0.2297 |
| CNAG_03206 | NTG2 | DNA-(apurinic or apyrimidinic site) lyase | 0.2266 | 3.5026 | 1.8853 | 0.4172 |
| CNAG_02315 | RIP1 | ubiquinol-cytochrome c reductase iron- sulfur subunit | 0.2213 | 2.4742 | 1.3029 | 0.4163 |
| CNAG_07667 | SAT4 | other/HAL protein kinase | 0.8944 | 0.9249 | 0.8341 | 0.9829 |
| CNAG_07356 | SHH3 | succinate dehydrogenase | 0.1641 | 2.1933 | 1.1275 | 0.3163 |
| CNAG_06558 | TAH18 | NADPH-ferrihemoprotein reductase | 0.8052 | 3.2042 | 1.8194 | 1.4057 |
| **Electron transport** |  |  |  |  |  |  |
| CNAG_05318 | CYB2 | L-mandelate dehydrogenase | 0.4106 | 2.7764 | 1.8402 | 0.6139 |
| CNAG_05169 | CYB2 | cytochrome b2 | 0.2067 | 4.5389 | 1.8602 | 0.4997 |
| CNAG_07480 | MCR1 | cytochrome b5 reductase | 2.2450 | 0.9528 | 1.6701 | 1.2686 |
| CNAG_00716 | CYC7 | electron carrier | 0.2159 | 2.1379 | 1.9578 | 0.2335 |
| CNAG_03666 |  | acyl-CoA dehydrogenase | 0.0559 | 2.8344 | 1.1617 | 0.1351 |
| CNAG_03629 | YHB1 | NADH-ubiquinone oxidoreductase | 0.0533 | 4.2998 | 1.1451 | 0.1982 |
| CNAG_01229 | CYB2 | L-mandelate dehydrogenase | 0.1334 | 3.9091 | 2.0502 | 0.2521 |
| CNAG_05631 |  | NADH-ubiquinone oxidoreductase | 0.0723 | 5.3982 | 1.2922 | 0.2991 |
| CNAG_01078 | ALD5 | aldehyde dehydrogenase | 1.5127 | 2.3937 | 2.9968 | 1.1969 |
| CNAG_04189 | SDH1 | succinate dehydrogenase flavoprotein subunit | 0.1650 | 1.5232 | 1.0283 | 0.2422 |
| CNAG_04521 |  | oxidoreductase | 1.4009 | 1.8313 | 2.5354 | 1.0024 |
| CNAG_03663 | CYB2 | L-lactate dehydrogenase | 0.1027 | 3.0248 | 1.3071 | 0.2355 |
| CNAG_03226 | SDH2 | succinate dehydrogenase iron-sulfur subunit | 0.0885 | 1.4843 | 0.8700 | 0.1495 |
| CNAG_05179 | QCR2 | ubiquinol-cytochrome C reductase complex core | 0.1674 | 2.1182 | 0.9486 | 0.3705 |
| CNAG_05258 |  | glucose-methanol-choline oxidoreductase | 5.6800 | 4.7204 | 17.3553 | 1.5288 |
| CNAG_01323 | QCR7 | ubiquinol-cytochrome-c reductase | 0.2675 | 3.1955 | 1.9649 | 0.4308 |
| CNAG_05909 | CYT1 | electron transporter | 0.1198 | 3.1845 | 1.1843 | 0.3193 |
| Mitochondria functions |  |  |  |  |  |  |
| CNAG_00162 |  | alternative oxidase | 0.0234 | 52.5633 | 1.7976 | 0.6774 |
| CNAG_03824 | Mir1 | phosphate transport protein MIR1 | 1.1861 | 0.2607 | 0.3061 | 1.0013 |
| CNAG_00499 |  | carnitine/acyl carnitine carrier | 0.3650 | 1.7017 | 1.0367 | 0.5939 |
| CNAG_05909 |  | electron transporter | 0.1198 | 3.1845 | 1.1843 | 0.3193 |
| CNAG_02315 |  | ubiquinol-cytochrome c reductase | 0.2213 | 2.4742 | 1.3029 | 0.4163 |
| CNAG_05132 |  | cytochrome c oxidase | 0.4774 | 1.1400 | 0.9550 | 0.5645 |
| CNAG_03225 |  | malate dehydrogenase | 0.9064 | 0.5619 | 0.4410 | 1.1443 |
| CNAG_00237 |  | 3-isopropylmalate dehydratase | 0.2215 | 9.5446 | 3.0096 | 0.6958 |
| CNAG_03596 |  | dihydrolipoamide succinyltransferase | 0.4599 | 0.8710 | 0.7416 | 0.5355 |
| CNAG_07908 |  | aconitate hydratase | 0.2025 | 5.6433 | 2.1318 | 0.5310 |
| CNAG_05031 |  | succinyl-CoA:3-ketoacid-coenzyme A transferase | 0.9592 | 0.2183 | 0.6530 | 0.3179 |
| CNAG_06138 |  | NADH dehydrogenase (ubiquinone) Fe-S 6 | 0.2215 | 2.0515 | 1.0108 | 0.4452 |
| DNA repair |  |  |  |  |  |  |
| CNAG_07552 | Rad8 | DNA repair Rad8 | 0.8674 | 6.7453 | 4.9744 | 1.1664 |
| CNAG_00178 | Rev1 | DNA repair REV1 | 1.4024 | 6.1169 | 7.1227 | 1.1940 |
| CNAG_05198 | Rad7 | DNA repair RAD7 | 1.2712 | 5.4930 | 7.2702 | 0.9515 |
| CNAG_02544 | Swi5 | DNA repair Swi5 Sae3 | 1.5561 | 5.0624 | 7.0387 | 1.1043 |
| CNAG_02512 | Rad16 | DNA repair RAD16 | 1.9665 | 4.8413 | 8.8648 | 1.0644 |
| CNAG_01163 | Rad54 | DNA repair and recombination RAD54 | 1.2403 | 3.4198 | 3.9817 | 1.0558 |
| CNAG_00299 | Rad54 | DNA repair RAD5 | 0.7534 | 2.8207 | 2.1946 | 0.9596 |
| CNAG_02771 | Rad54b | DNA repair and recombination RAD54B | 1.0984 | 7.9051 | 8.7877 | 0.9793 |
| CNAG_00720 | Rad51 | DNA repair RAD51 | 1.5080 | 7.1115 | 10.7208 | 0.9912 |
| CNAG_03637 |  | Double strand break repair factor and silencing regulator | 1.0924 | 3.6090 | 3.7124 | 1.0530 |
| Ergosterol metabolism |  |  |  |  |  |  |
| CNAG_00519 | ERG3 |  | 2.1758 | 0.2356 | 0.3907 | 1.3001 |
| CNAG_01129 | ERG7 |  | 1.6644 | 0.4326 | 0.5393 | 1.3235 |
| CNAG_00854 | ERG2 |  | 2.4636 | 0.2523 | 0.5729 | 1.0753 |
| CNAG_02830 | ERG4 |  | 1.8153 | 0.4589 | 0.9389 | 0.8795 |
| CNAG_01737 | ERG25 |  | 2.1911 | 0.1713 | 0.3510 | 1.0601 |
| Oxidative stress |  |  |  |  |  |  |
| CNAG_00575 | Cat3 | catalase 3 (CAT3) | 0.4810 | 5.5936 | 3.3113 | 0.8059 |
| CNAG_00654 | Srx1 | conserved hypothetical protein (SRX1, sulfiredoxin) | 4.3449 | 4.8477 | 14.4082 | 1.4488 |
| CNAG_01138 | Ccp1 | cytochrome c peroxidase (CCP1) | 0.0674 | 181.0047 | 27.3521 | 0.4416 |
| CNAG_03482 | Tsa1 | thioredoxin-dependent peroxide reductase (TSA1) | 1.3127 | 5.0984 | 4.6206 | 1.4351 |
| CNAG_03985 | Grx5 | monothiol glutaredoxin-5 (GRX5) | 0.4517 | 1.9536 | 2.0371 | 0.4293 |
| CNAG_04388 | Sod2 | mitochondrial superoxide dismutase Sod2 | 0.9830 | 0.5836 | 0.5057 | 1.1240 |
| CNAG_05847 | Trr1 | thioredoxin-disulfide reductase (TRR1) | 1.2419 | 12.1215 | 10.1329 | 1.4726 |

The numbers in pink are those measurements where the log2 value is greater than or equal to 2; the numbers in green are log2 values of less than 0.5.

We next constructed heat maps of specific functions highlighted by the GO term analysis to further examine the influence of Grx4 on transcription. In the context of GO terms for molecular function, loss of Grx4 influenced transcript levels for metal ion transport (Fig. 8A), iron sulfur cluster binding (Fig. 8B) and heme biosynthesis (Fig. 8C), as shown. Loss of Grx4 generally resulted in elevated transcripts for these genes, a finding consistent in part with the participation of Grx4 in the regulation of iron-using functions. We also constructed heat maps to examine the expression patterns of genes in additional categories from the GO term analysis and these revealed further connections between Grx4 an iron-using functions for electron carrier activity (Fig. 9). As noted above, DNA repair, replication and recombination functions were the top category for the GO term analysis for biological processes. The expression patterns for genes in this category are shown in Fig. 10, and this regulation is consistent with the important role of iron as a cofactor for enzymes involved in DNA synthesis and repair (41, 42).

**Figure 8.**
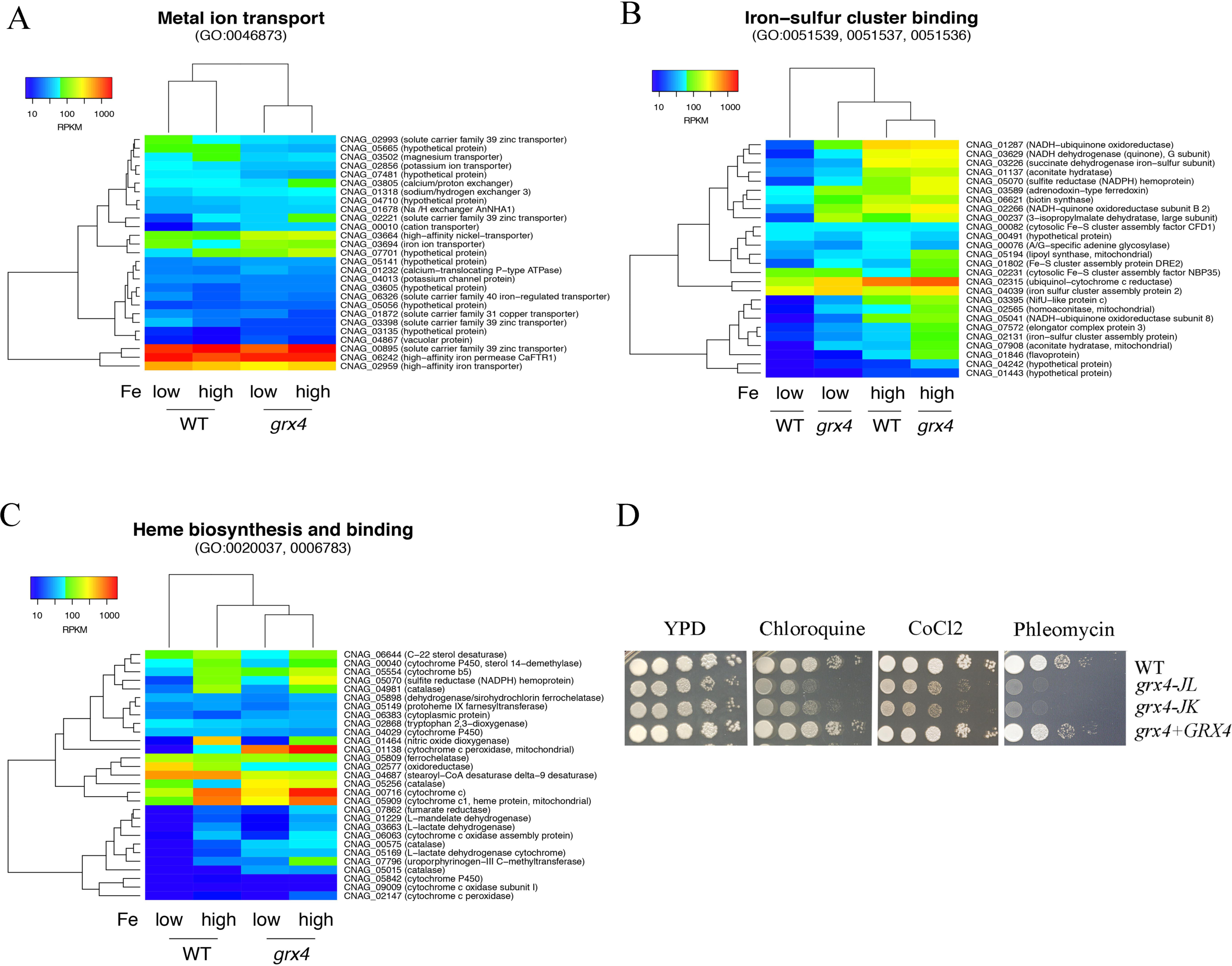
Grx4 regulates genes involved in metal ion transport, heme biosynthesis and iron-sulfur cluster binding in response to iron availability. A) Changes in transcript abundance of the genes encoding functions in metal ion transport between the WT and *grx4* mutant strains grown in low iron and high iron conditions and the results were represented by the heat-map. B) Changes in transcript abundance of the genes encoding functions in iron-sulfur cluster binding between the WT and *grx4* mutant strains grown in low iron and high iron conditions and the results were represented by the heat-map. C) Changes in transcript abundance of the genes encoding functions in heme biosynthesis and binding between the WT and *grx4* mutant strains grown in low iron and high iron conditions and the results were represented by the heat-map. D) Tenfold serial dilutions of each strain, including the WT, two independent *grx4* disruption mutants, the *GRX4* complemented strains, without iron starvation were spotted on the YPD plates with or without the antimalarial drug chloroquine (6 mM), CoCl_2_ (a hypoxia-mimicking agent, 600 μM) or phleomycin (an iron-dependent inhibitor, 8 pg/ml). The plates were incubated at 30 °C for 2 days before being photographed.

#### Loss of Grx4 results in phenotypes consistent with dysregulation of iron homeostasis

The transcriptome changes in both low- and high-iron conditions (Fig. 7) strongly indicate that iron homeostasis is dysregulated upon loss of Grx4. We therefore examined iron-related phenotypes to confirm the biological impact of the Grx4 defect. Specifically, we tested the sensitivity of the *grx4* mutants to a wide range of inhibitors that influence iron-dependent processes including chloroquine, the hypoxia-mimicking agent CoCl_2_, and the iron-dependent inhibitor phleomycin (Fig. 8D). Additionally, our phenotypic assays with inhibitors of complexes of the electron transport chain and agents that provoke oxidative stress demonstrated sensitivity for the *grx4* mutants (Fig. 9B-D). Finally, we found that the *grx4* mutants were sensitive to agents that provoke oxidative stress or DNA damage (Fig. 10B). The phenotypes indicated in Figure. 8–10 were observed both on rich medium (as shown) and on the more defined YNB medium supplemented with iron (Supplemental Fig. S7). Overall, the observed phenotypes are consistent with a role for Grx4 in regulating iron homeostasis such that iron-dependent functions are dysregulated upon loss of the protein. These functions are likely to be critical for mitochondrial function and adaptation to the host environment.

**Figure 9.**
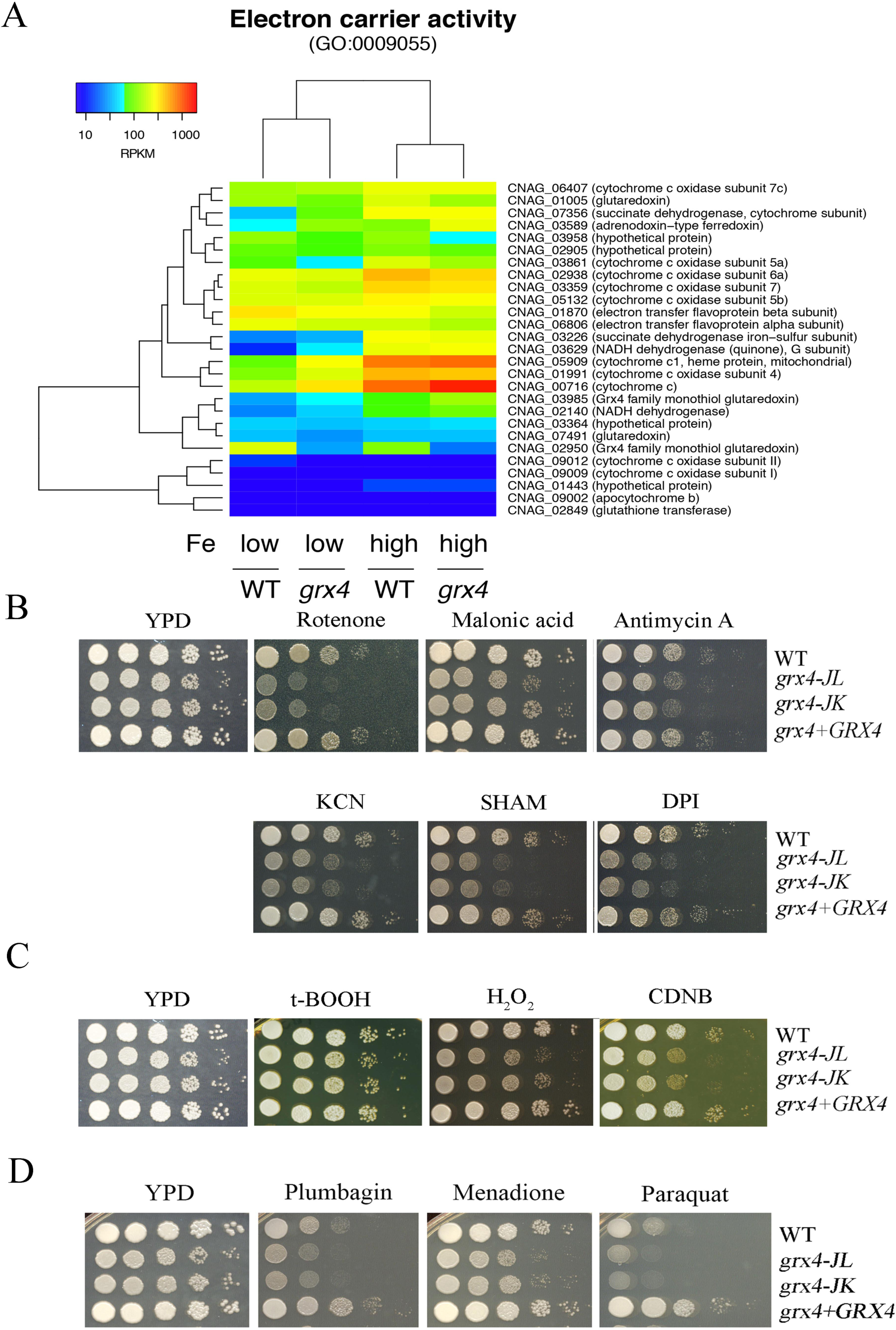
Grx4 is implicated in the regulation of functions for electron transport and the response to oxidative stress. A) Changes in transcript abundance of the genes encoding functions in electron carrier activity between the WT and *grx4* mutant strains grown in low iron and high iron conditions and the results were represented by the heat-map. B) Disruption of GRX4 leads to sensitivity to inhibitors of electron transport chain complexes I to IV and the alternative oxidase (75 μg/ml rotenone, 2 mM malonic acid, 5 μg/ml antimycin A, 10 mM potassium cyanide (KCN), 10 mM salicylic hydroxamate (SHAM) and 50 μM diphenyleneiodonium (DPI)). Tenfold serial dilutions of each strain were spotted onto the YPD plates supplemented with the drug indicated and the plates were incubated at 30°C for 2 days before being photographed. C) Disruption of GRX4 leads to the sensitivty to some agents that provoke oxidative stress (2mM t-BOOH, 0.01% H_2_O_2_ and 5 μg/ml 1-chloro-2,4-dinitrobenzene (CDNB)). Tenfold serial dilutions of each strain were spotted onto the YPD plates supplemented with the drug indicated and the plates were incubated at 30°C for 2 days before photographed. D) The grx4 mutants have altered susceptibilities to inhibitors of reactive oxygen species (50 μM plumbagin, 5 μg/ml menadione, and 500 μM paraquat). Tenfold serial dilutions of each strain were spotted onto the YPD plates supplemented with the drug indicated and the plates were incubated at 30°C for 2 days before photographed. Plates were incubated at 30°C and 37°C for 2 days.

**Figure 10.**
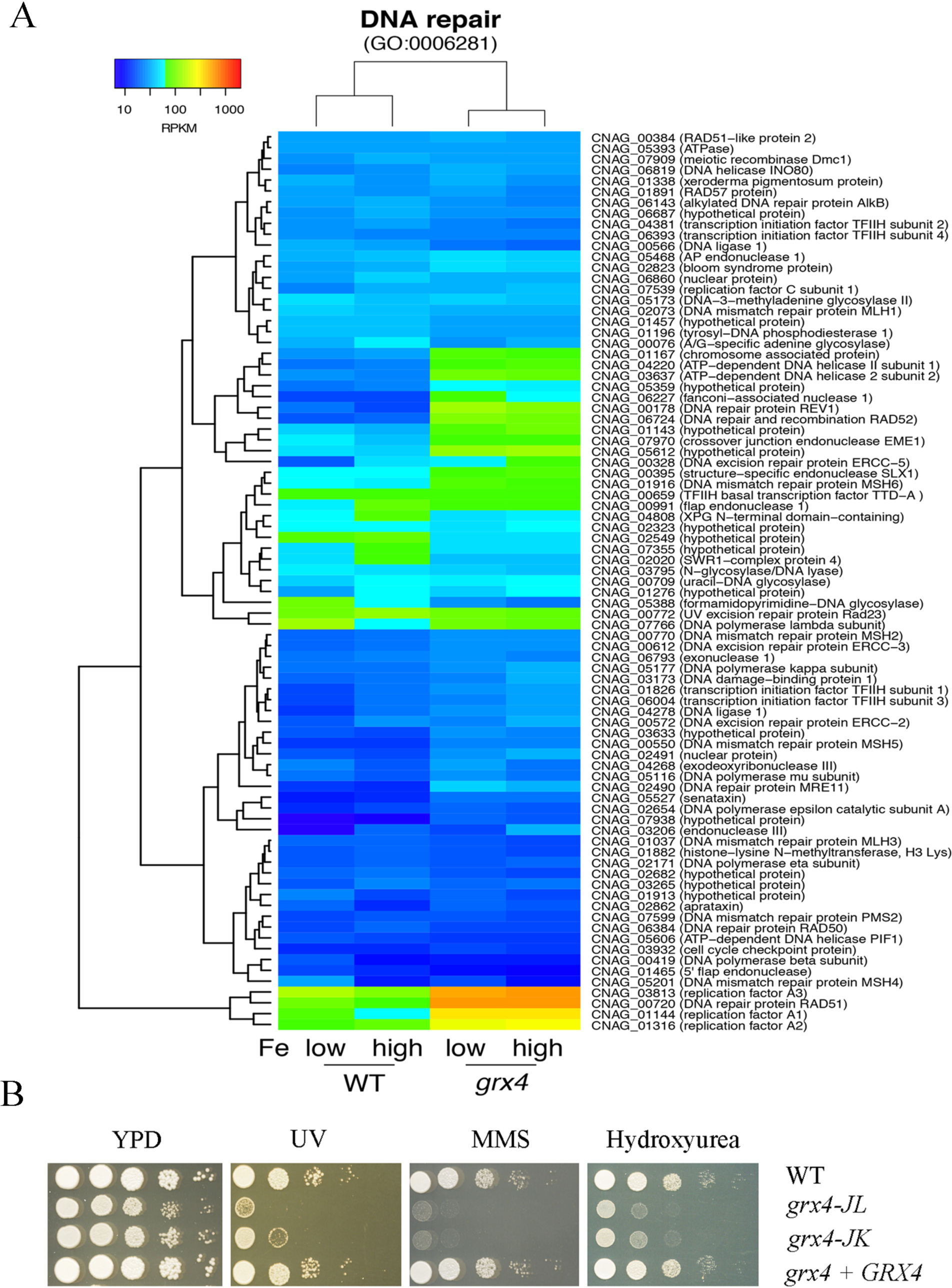
Grx4 regulates function for DNA repair and confers resistance to DNA damaging agents. A) Changes in transcript abundance of the genes encoding functions in DNA repair between the WT and *grx4* mutant strains grown in low iron and high iron conditions and the results were represented by the heat-map. B) Tenfold serial dilutions of each strain were spotted onto the YPD plates supplemented with the drugs indicated and the plates were incubated at 30°C for 2 days before being photographed. In addition to exposure to UV light (100 and 400 J/m2), the drugs included DNA repair inhibitors 25 mM hydroxyurea (HU) and 0.03% methyl methanesulfonate (MMS).

## Discussion

In this study, we identified a putative monothiol glutaredoxin Grx4 as an interaction partner with Cir1, the iron-responsive transcription factor that regulates iron uptake functions and virulence in *C. neoformans.* Subcellular localization studies reinforced the idea that interaction between Grx4 and Cir1 is relevant for iron sensing. That is, we found that Grx4 moves from the nucleus to the cytoplasm upon iron repletion, while Cir1 is located in the nucleus regardless of iron availability. Interestingly, the relocation of Grx4 was dependent on Cir1 because a Grx4-mCherry fusion protein remains in the cytoplasm in a *cir1* mutant. Additionally, more than one factor may contribute to retention of Grx4 in the nucleus given that treatment with the proteasome inhibitor BTZ influenced the location of the Grx4-mCherry signal. These findings prompted a detailed characterization of the impact of a *grx4* mutation on iron homeostasis and virulence. We found that Grx4 is required for robust proliferation upon iron depletion and at 37^°^C, and for the elaboration of major virulence factors including capsule and melanin. These phenotypes were consistent with an observed virulence defect in a murine inhalation model of cryptococcosis. Subsequent transcriptional profiling revealed that Grx4 influences that expression of genes for a variety of iron-dependent functions including DNA repair, response to oxidative stress, [2Fe-2S] cluster binding, heme binding and oxidoreductase activity. Consistent with this regulation, a *grx4* mutant showed increase sensitivity to agents such as inhibitors of electron transport complexes that challenge functions that utilize iron.

The wealth of information from model yeasts on the mechanistic details of monothiol GRXs, [2Fe-2S] cluster binding and transcriptional regulation provides a framework to interpret the contribution of Grx4 to iron sensing in *C. neoformans* (25–34, 43). In model yeasts, monothiol glutaredoxins play a critical role in sensing iron availability via [2Fe-2S] cluster assembly to influence the activities of transcription factors that regulate the expression of iron acquisition and iron dependent functions (25–34). For example, the monothiol glutaredoxins Grx3 and Grx4 form [2Fe-2S] cluster binding complexes with the cytosolic proteins Fra1 and Fra2 in *S. cerevisiae* to dissociate the activator Aft1 from the promoters of genes for iron uptake upon iron repletion. As a result Aft1 is relocated from the nucleus to the cytoplasm. In contrast, iron deprivation results in accumulation of Aft1 in the nucleus where it activates iron uptake, mobilization of stored iron from the vacuole and remodeling of iron-dependent metabolism (44). As mentioned, Grx4 regulates the iron-responsive transcription factors Fep1 and Php4 in *S. pombe.* Under iron-replete conditions, Grx4 binds and inactivates Php4, a repressor of genes encoding proteins for iron use. In this condition, Php4 is retained in the cytoplasm in a Grx4-dependent manner (39). Upon iron limitation, the association of Grx4 and Php4 is reduced, and Php4 accumulates in the nucleus to repress genes encoding proteins for iron use. Deletion of Grx4 makes Php4 constitutively active and permanently located in the nucleus. Fep1 is an iron-containing protein, and bound iron is required for transcriptional repression of iron uptake functions in iron-replete conditions (24, 27, 29, 30). A Grx4-Fra2 heterodimer constitutively binds to Fep1 and iron deprivation results in disassembly of the Fe-S cluster between Grx4 and Fra2 to allow metal transfer from Fep1 to Grx4-Fra2 and derepression of iron uptake functions.

Our analysis revealed novel features of Grx4 and Cir1 compared with the studies in the model yeasts. Comparisons are most informative with *S. pombe* because of the detailed information on Fep1, a candidate ortholog of Cir1 in *C. neoformans.* The interaction of Grx4 with Cir1 resembles that of Grx4 with Fep1, although Cir1 has a broader transcriptional impact beyond the regulation of iron uptake functions (15, 16). However, we don’t yet have the same detailed molecular understanding of the impact of Grx4 on Cir1 activity that is available for Fep1 in *S. pombe* (26, 27, 29). One distinction for *C. neoformans* is the finding that the nuclear versus cytoplasmic location of the Grx4 protein is responsive to iron and heme. This is not the case for Grx4 *in S. pombe* where the protein is located in both compartments regardless of iron availability (29). As mentioned above, the Grx4 partner Php4 relocates between the nucleus and the cytoplasm in response to iron repletion in *S. pombe*. Php4 is simlar to Hap4 in *S. cerevisiae* and is part of the well-characterized CCAAT complex (Hap2, 3, 4, 5) (45). Orthologs in *Candida albicans* (Hap43) and *Aspergillus fumigatus* (HapX) are also involved in iron regulation and virulence (46–48). We previously characterized the function of a Php4-related protein designated HapX in *C. neoformans* (16). Microarray analysis revealed that HapX influences transcript levels of genes encoding iron use functions, as expected for an ortholog of Php4. HapX may therefore be an additional partner of Grx4 in *C. neoformans.* Interestingly, HapX positively regulates the transcript levels of *CIR1* and genes encoding iron uptake functions in the low-iron condition. Like Php4, HapX also negatively regulates functions that use iron such as electron transport proteins. Given that Grx4 and Php4 interact in *S. pombe*, we predict that Grx4 and HapX proteins have similar interactions and regulatory functions in *C. neoformans.* Experiments to examine this prediction are currently underway, and our preliminary yeast two-hybrid assays indicate that Grx4 interacts with HapX (E. Sanchez-Leon, unpublished results). In this context, it is interesting to note that Grx4 makes a greater contribution to the ability of *C. neoformans* to cause disease than HapX (16) suggesting a wider influence on functions that contribute to proliferation in vertebrate hosts.

The interaction of Grx4 with transcription factors in model yeasts suggests that Grx4 may co-regulate genes with Cir1 in *C. neoformans*. We found that loss of Grx4 impacted the expression of a partially overlapping set of genes compared with a previous microarray study in which we identified the sets of genes regulated by Cir1 and HapX in response to different iron levels (15, 16). The GO terms for Cir1 regulation under low iron included iron ion transport, siderophore transport, processing of 20S pre-RNA, and rRNA metabolism (16). Additional GO term categories of DNA replication, DNA metabolism, and DNA repair were also identified in an earlier study (15). For HapX, we found that loss of this factor influenced the transcript levels for genes in the GO categories of ATP synthesis coupled electron transport, and siderophore transport in the low-iron condition (16). As expected, Grx4 regulates a subset of genes in the categories influenced by Cir1 and HapX. Specifically, the GO categories for transcripts impacted by the loss of Grx4 included double-strand break repair, response to oxidative stress and oxidation-reduction processes, respiratory electron transport chain, energy derivation by oxidation of organic compounds and carbohydrate derivative metabolic process. In this regard, the pattern for Grx4 more closely resembles that of HapX, especially for functions related to electron transport. We therefore speculate that part of the contribution of Grx4 occurs through an interaction with HapX, a protein with similarities to Php4 in *S. pombe*, and a shared contribution to the regulation of iron-using functions. Given that only microarray data is currently available for Cir1 and HapX, additional work is needed to obtain RNA-Seq data for mutants lacking these proteins to allow a more direct comparison of shared and distinct targets of regulation with Grx4.

We confirmed the RNA-Seq finding that Cir1 and Grx4 both participate in the regulation of a subset of genes by examining the targets *LAC1* and *FRE3* with qRT-PCR. Loss of Grx4 or Cir1 results in elevated *LAC1* transcripts but the impact melanin formation is quite different. That is, a *cir1* mutant causes a hyper-melanized phenotype and our current analysis revealed a melanin defect for the *grx4* mutant (15). These observations suggest that Grx4 might influence the expression of *LAC1* by an additional mechanism that is independent of Cir1, and in these regard it is likely that Grx4 may interact with other transcription factors (in addition to HapX). Some of these other factors may influence melanization. Media conditions could also have an influence because the cells for RNASeq analysis were grown in liquid media and L-DOPA solid medium was used to assay melanin. Cir1 and Grx4 also positively regulated *FRE3*, as expected for shared participation in the control of a subset of iron uptake functions.

Why does loss of Grx4 cause a severe virulence defect? A significant component of the contribution of Grx4 is likely due to its influence on the expression and activity of Cir1, as well as potential regulatory interactions with other transcription factors such as HapX. In particular, the *grx4* and *cir1* mutants share defects in capsule formation, the major virulence factor, and both fail to proliferate well at 37^°^C. These phenotypes would certainly be expected to impair virulence. Other contributions are likely and these include the dysregulation of iron homeostasis in the *grx4* mutant to impair adaptation to the host environment necessary to withstand defense responses (e.g., oxidative stress). In this context, human glutaredoxin is known to play an important role in redox homeostasis and protection against oxidative damage (49). Similarly, Grx3 and Grx4 in *S. cerevisiae*, Grx3 in *C. albicans*, and Grx3 in the insect pathogen *Beauveria bassiana* contribute to resistance to oxidative stress (50–53). A complete understanding of the contribution of Grx4 will require future work on the functional implications of interactions with Cir1, HapX and other transcription factors, and an investigation of the mechanisms of iron sensing in host tissue.

## Materials and Methods

### Strains, plasmids, chemicals, and media

The serotype A strain H99 (MATα) of *C. neoformans* var. *grubii* and mutant derivatives were maintained on YPD medium (1% yeast extract, 2% peptone, 2% dextrose, 2% agar). The nourseothricin, neomycin, and hygromycin resistance cassettes were from plasmids pCH233, pJAF1, and pJAF15 (obtained from Dr. J. Heitman), respectively. YPD medium plates containing nourseothricin (100 μg/ml) were used to select the *grx4* deletion transformants, and plates containing neomycin (200 μg/ml) to select the *GRX4* reconstituted transformants from the *grx4* background. Defined low-iron medium (LIM) and yeast nitrogen base (YNB with amino acids; pH 7.0) plus 150 μM bathophenanthroline disulfonate (BPS) (YNB-LIM) were used as iron-limiting media. YPD and/or YNB media supplemented as indicated were used for phenotypic characterizations in growth assays on liquid and solid media.

### Capsule formation and melanin production

Capsule formation was examined by differential interference contrast microscopy after incubation for 24 h to 48 h at 30°C in defined LIM and staining with India ink. Melanin production was examined on L-3,4-dihydroxyphenylalanine (L-DOPA) plates containing 0.1% glucose.

### Protein-protein interaction assays

The Grx4-Cir1 interaction assays were performed using the ProQuest Two hybrid system with Gateway Technology, Invitrogen Life Technologies Inc., according to the manufacturer’s protocols, and as previously described (37). Briefly, the coding sequence for *GRX4* (726 bp) was synthesized by BioBasic (www.biobasic.com/gene-splash) and cloned into pDEST-22 (Gal4 DNA binding domain). The N-terminal region of *CIR1* (1-1203 bp) was PCR amplified from *C. neoformans* cDNA using the primers listed in Supplemental Table S4, and cloned into pDEST-32 (Gal4 activating domain). The plasmids were then co-transformed into MaV203 yeast competent cells.

The growth of MaV203 yeast expressing both bait and prey vectors was tested on synthetic complete medium (0.7% Yeast nitrogen base without amino acids (Difco), 2.0% glucose, 0.07% synthetic complete selection medium mix (Sigma), 1.7% bacto agar (Difco), pH 5.6) lacking leucine and tryptophan to select for each vector, and histidine and uracil to test for an interaction. Empty pDEST-32 and pDEST-22 vectors were used as negative controls. The physical interaction between the encoded proteins in these plasmids was tested by assessing restoration of uracil and histidine prototrophy, and by assaying the activity of β-galactosidase (according to manufacturer’s protocols, and as previously described (37)).

### Deletion of the conserved Grx domain to create a *grx4* mutant and generation of *grx4Δ::GRX4* complemented strain

A *grx4* disruption mutant was constructed by homologous recombination using a nourseothricin acetyltransferase (NAT) marker linked to 5’ and 3’ flanking sequences of *GRX4* >C-terminal domain by three-step overlapping PCR using primers listed in Supplemental Table S4. The overlap PCR product was biolistically transformed into the WT strain H99, and deletion was confirmed by PCR and Southern blot hybridization as previously described (54–56). Genomic DNA for Southern blot analysis was prepared using cetyl trimethyl ammonium bromide (CTAB) phenol-chloroform extraction.

To reconstitute the deleted region of *GRX4* in the *grx4* mutant, a genomic DNA fragment containing 1.1 kb of upstream promoter region and a 1.7 kb region carrying the deleted portion of *GRX4* gene was amplified by PCR. This PCR fragment was fused with the neomycin (NEO^r^) selectable marker (1.9 Kb) at its C terminus in an overlap PCR reaction. The overlap PCR product was introduced into the *grx4* mutant by biolistic transformation. Targeted integration was confirmed by PCR and genomic hybridization.

### Construction of a *CIR1::GFP* fusion allele

The C-terminal region of the Cir1 protein was tagged with GFP (green fluorescent protein) to examine the subcellular localization of Cir1. Briefly, the left arm (836 bp) and right arm (826 bp) for the fusion construct were amplified from WT gDNA using the primer set Cir1-GFP-P1F and Cir1-GFP-P1R and the primer set Cir1-GFP-P5F and Cir1-GFP-P5R, respectively. *GFP* and the hygromycin (*HYG*) resistance gene were amplified from the plasmid pGH022 using primers Cir1-GFP-P2F and Cir1-GFP-P3R (3,476 bp). Overlap PCR was performed using primers Cir1-GFP-P1F and Cir1-GFP-P5R to yield the 5,138 bp construct. The construct was then used to transform the WT and *grx4* mutant strains by biolistic transformation. Following transformation, mutants were screened for resistance to HYG and proper location and orientation of *GFP* was determined by PCR. Primer sequences are listed in Table S4.

### Construction of a *GRX4::mCherry* fusion allele

The C-terminal region of the Grx4 protein was tagged with mCherry to examine the subcellular localization of Grx4. Briefly, the left arm (548 bp) and right arm (508 bp) for the fusion construct were amplified from WT gDNA using the primer set Grx4-mCherry-P1F and Grx4-mCherry-P1R and the primer set Grx4-mCherry-P3F and Grx4-mCherry-P3R, respectively. The gene encoding mCherry and the hygromycin (*HYG*) resistance gene were amplified from the plasmid pGH026 using primers Grx4-mCherry-P2F and Grx4-mCherry-P2R (3,524 bp). Overlap PCR was performed using primers Cir1-GFP-P1F and Cir1-GFP-P3R to yield the 4,580 bp construct. The construct was then used to transform the WT (H99), WT::Cir1-GFP, and *cir1* mutant strains by biolistic transformation. Following biolistic transformation, mutants were screened for resistance to HYG and the proper location and orientation of mCherry was determined by PCR. Primer sequences are listed in Table S4.

### Quantitation of Grx4-mCherry fluorescence

Laser scanning confocal fluorescence images were analyzed to determine signal intensities of mCherry fluorescence in the cytoplasm and nuclei in response to different iron sources/levels for the *C. neoformans* strain co-expressing Grx4-mCherry and Cir1-GFP. The mean gray values of selected fluorescent areas (Integrated Density) showing Grx4-mCherry corresponding to the nucleus and the whole cell were obtained for each individual cell. Nonnuclear fluorescent signals were determined by the subtraction of Integrated Density of the whole cell (WC) and the nucleus (N). All measurements were corrected for each individual image background. The scatter plot graph shows each treatment mean with confidence intervals (CI) 95% (n≥25). One-way ANOVA and Tukey statistical tests were performed to analyze the signal intensity data of the *C. neoformans* strain under low iron (LIM) and iron supplemented (10μM FeCl_3_/Hemin) treatments. Both tests revealed statistical significant differences for the different treatments *(p* < 0.0001). Laser scanning confocal fluorescence microscopy was performed with an inverted Olympus Fluoview FV1000 confocal microscope fitted with an argon laser (GFP: excitation, 488 nm; emission, 510 nm) and a He/Ne laser (mCherry: excitation, 543 nm; emission, 612 nm); along with a 100× UplanS Apo (Olympus) oil-immersion objective (NA, 1.40). Confocal images were captured and examined using FV10-ASW software (version 4.2.3.6, Olympus). Fluorescence signal intensity measurements and statistical analysis were performed with ImageJ (version 1.51w; Maryland, USA) and Prism 6 (version 6.01; GraphPad Software).

### RNAseq analysis of gene expression

Cells for three biological replicates were prepared by growing the WT strain and *grx4*Δ mutant in 50 ml of YPD for overnight at 30°C. The cells were washed twice with iron-chelated water followed by growth in 50 ml of YNB low-iron medium for 16 hours, harvested and diluted to 4.0 × 107 cells in 50 ml of the same medium with or without 100 μM of FeCl3. Cultures were incubated at 30°C for 6 hours and harvested for RNA extraction. Total RNA was extracted by RiboPure-Yeast Kit (Life Technologies, Carlsbad, CA, USA) and treated DNase I (Life Technologies, Carlsbad, CA, USA) following the manufacturer’s instructions. The quantity and integrity of the total RNA were evaluated using 2100 Bioanalyzer (Agilent Technologies, Palo Alto, CA, USA). The samples for verification of RNAseq data by qPCR were prepared as the same as above. The primers used for qPCR are listed in Supplemental Table S5.

### Read alignment and quantification

Raw Illumina reads were quality-trimmed and filtered for adapter contamination using Trimmomatic v. 0.36 (57). The following settings were applied: “ILLUMINACLIP:TruSeq3-PE.fa:2:30:10 LEADING:10 TRAILING:10 SLIDINGWINDOW:5:10 MINLEN:50”. Reads passing the filters were aligned to the latest *C. neoformans* var. *grubii* H99 reference genome using hisat v. 2.1.0 (58) with parameters –min-intronlen 20 –max-intronlen 1000. The genome sequence and annotation were downloaded from ENSEMBL (http://fungi.ensembl.org) in August 2017. The produced SAM files were position-sorted using samtools v. 1.5 (Li *et al.*, 2009). Reads overlapping annotated transcripts of H99 were quantified with HTSeq-count v. 0.9.1 requiring a minimum alignment quality of 10 and setting the matching mode to “union” (59).

Read counts were normalized across samples and replicates using the Biocondutor package edgeR v. 3.6 (61). We removed genes with total counts below 3 prior to normalization (62). Library sizes were normalized using Trimmed Mean of M values (TMM) and gene-level counts were normalized across conditions using RPKM (Reads per kilobase of transcript per million mapped reads). For this, the edgeR functions *cpm* and *rpkm* were used. Differential expression was tested using the edgeR function *exactTest*, which tests for mean expression differences based on negative-binomially distributed counts. Significance values were corrected for multiple testing using the Benjamin-Hochberg False Discovery Rate (FDR). We restricted our analyses to genes with an expression difference of at least two-fold and a FDR *p*-value < 0.001. Log10-scaled gene expression values were visualized using the *heatmap.2* function in the R package gplots v. 3.0.1.

### Analyses of enrichment in protein functions

For each comparison among conditions (iron levels and *grx4* genotypes), sets of differentially expressed genes were analyzed for an enrichment in encoded protein functions. Predicted proteins were assigned to Gene Ontology (GO) terms using InterProScan 5.26-65 (63). GO terms were only considered if the total term size in the genome was at least 5. For each comparison, hypergeomtric tests were performed to test for enrichment and GO terms with a FDR *p*-value < 0.001 were considered as significant. All enrichment analyses were performed using the R packages GSEABase and GOstats (64). Outcomes of enrichment tests were visualized using the R package ggplot2 (65).

### Virulence assay

For virulence assays, female BALB/c mice (4–6 weeks old) were obtained from Charles River Laboratories (Ontario, Canada). The WT, *grx4* mutant and *grx4Δ::GRX*cells were grown in YPD overnight at 30°C, washed in PBS and re-suspended at 1.0 × 10^6^ cells ml^−1^ in PBS. Inoculation was by intranasal instillation with 50 μl of cell suspension (inoculum of 2.0 × 10^5^ cells per mouse). Groups of 10 mice were inoculated for each strain. The status of the mice was monitored twice, daily post inoculation. For the determination of fungal burdens in organs, infected mice were euthanized by CO_2_ inhalation and organs were excised, weighed and homogenized in 1 ml of PBS using a MixerMill (Retsch). Serial dilutions of the homogenates were plated on YPD agar plates containing 35 Δg ml^−1^ chloramphenicol and CFUs were counted after incubation for 48 h at 30°C.

Furthermore, fungal load distribution in different tissues of the infected mice was determined. Mice reaching the endpoint were euthanized by CO_2_ asphyxiation, and fungal loads in different tissues of the mice including the lungs, kidney, liver, spleen, brain and blood were determined. Tissues were aseptically removed and immersed in PBS. Organs were homogenized using an automated tissue homogenizer. The samples were serially diluted in PBS and plated on YPD supplemented with 35 μg/mL chloramphenicol. After two days of incubation at 30° C, the colony forming units (CFUs) were counted manually. All experiments with mice were conducted in accordance with the guidelines of the Canadian Council on Animal Care and approved by the University of British Columbia’s Committee on Animal Care (protocol A17-0117).

## Acknowledgements

We thank Debbie Adam and the Biodiversity Research Centre at UBC for sequencing support, and Celine Chan for technical assistance. This work was supported by a grant (5R01 AI053721) from the National Institute of Allergy and Infectious Diseases (to JK), and a fellowship from the Canadian Institutes of Health Research (to RA). DC is supported by the Swiss National Science foundation grant 31003A_173265. JK is a Burroughs Wellcome Fund Scholar in Molecular Pathogenic Mycology.

## Supplemental material

**Figure S1. Grx4 from *C. neoformans* has a conserved C-terminal GRX domain with a signature “CGFS” motif.** A) Alignment of the amino acid sequence of the C-terminal Grx domain of *C. neoformans* Grx4 (XP_012047837.1) with selected Grx sequences from other organisms. UhGrx4, *Ustilago hordei*, CCF50760.1; UmGrx4: *Ustilago maydis*, XP_011390702.1; SpGrx4, *Schizosaccharomyces pombe*, NP_596647.1; ScGrx3, *Saccharomyces cerevisiae*, AJV06961.1; ScGrx4, *S. cerevisiae*, NP_011101.3; CaGrx4, *Candida albicans*, KHC50815.1; HsGrx4, *Histoplasma capsulatum*, EEH02725.1; HcGrx4, *Homo sapiens*, AAF28841.1. B) Phylogeny tree generated using the neighbouring joining tree analysis in MEGA(ver.7.0.26).

**Figure S2. The phenotypes of tagged strains (Grx4-mCherry, Cirl-GFP and Grx4-mCherry Cirl-GFP) are the same as the WT strain.** Tenfold serial dilutions of each strain grown in YPD medium overnight were spotted onto YPD or DOPA. The plates were incubated at either 30 °C or 37^°^C for 2 days before being photographed.

**Figure S3. Abundance of Cirl-GFP or Cirl-mCherry assessed by western blot analysis.** A) The extracted proteins from cells expressing Grx4-mCherry and WT cells grown under low iron condition were analyzed by Western blotting using anti-mCherry antibody. Intact Grx4-mCherry (58 kDa) was detected in Grx4-mCherry-expressing cells but not in WT cells. B) The extracted proteins from Grx4-mCherry-expressing cells in either WT or the *cir1* mutant background grown under the indicated conditions (low iron verse high iron) for 5 hours were analyzed by Western blotting using anti-mCherry antibody. C) The extracted proteins from Cir1-GFP-expressing cells in either the WT or *grx4* mutant background grown under indicated conditions (low iron verse high iron) for 5 hours were analyzed by Western blotting using anti-GFP antibody.

**Figure S4. Co-localization of Cirl-GFP with DAPI.** Cir1-GFP is localized in nuclei as detected by DAPI staining and independent of iron availability.

**Figure S5. STRING analysis of up-regulated transcripts in the *grx4* cells in either low or high iron condition.** Grx4 regulates the mitochondria relevant functions (cellular respiration, electron transport and metal binding), and oxidative phosphorylation. STRING was used to visualize predicted protein-protein interactions for the identified 639 Grx4-regulated proteins (http://string-db.org) using the corresponding proteins from *C. neoformans* strain JEC21 in the database.

**Figure S6. Verification of RNAseq data by q-PCR. Q-PCR was used to verify the expression of *CIR1, LAC1* and *FRE3*, in the WT and *grx4* mutant cells, grown in either low iron (A) or high iron (B) condition.** The expression pattern of these three genes is consistent with the data revealed by RNAseq analysis.

**Figure S7. Grx4 influences susceptibility to various inhibitors and DNA damaging agents on minimal medium supplemented with iron.** Tenfold serial dilutions of each strain were spotted onto YNB medium plus BPS and FeCl_3_ (100 μM) supplemented with the agents indicated, and the plates were incubated at 30°C for 2 days before being photographed. The concentrations of the inhibitors were as follows: chloroquine (6 mM), CoCl2 (600 μM), phleomycin (8 μg/ml), SHAM (10 mM), and paraquat (500 μM). DNA damaging agents included UV light (100 J/m2) and 0.03% methyl methanesulfonate (MMS).

Table S1. Comparison of Grx4 protein identities.

Table S2. List of transcripts up regulated in the *grx4* muant in both low and high iron conditions.

Table S3. List of transcripts down regulated in the *grx4* muant in both low and high iron conditions.

Table S4. Primers for strain construction.

Table S5. Primers for qRT-PCR.

## References

1. Park BJ, Wannemuehler KA, Marston BJ, Govender N, Pappas PG, Chiller TM. 2009. Estimation of the current global burden of cryptococcal meningitis among persons living with HIV/AIDS. AIDS 23:525–530.

2. May RC, Stone NR, Wiesner DL, Bicanic T, Nielsen K. 2016. *Cryptococcus:* from environmental saprophyte to global pathogen. Nat Rev Microbiol 14:106–117.

3. Ballou ER, Johnston SA. 2017. The cause and effect of Cryptococcus interactions with the host. Curr Op Microbiol 40:88–94

4. Rajasingham R, Smith RM, Park BJ, Jarvis JN, Govender NP, Chiller TM, Denning DW, Loyse A, Boulware DR. 2017. Global burden of disease of HIV-associated cryptococcal meningitis: an updated analysis. Lancet Infect Dis 17:873–881.

5. Jung WH, Kronstad JW. 2008. Iron and fungal pathogenesis: a case study with *Cryptococcus neoformans*. Cell Microbiol 10:277–284.

6. Kronstad JW, Attarian R, Cadieux B, Choi J, D’Souza CA, Griffiths EJ, Geddes JM, Hu G, Jung WH, Kretschmer M, Saikia S, Wang J. 2011. Expanding fungal pathogenesis: *Cryptococcus* breaks out of the opportunistic box. Nat Rev Microbiol 9:193–203.

7. Kronstad JW, Hu G, Jung WH. 2013. An encapsulation of iron homeostasis and virulence in Cryptococcus neoformans. Trends Microbiol 21:457–465.

8. Bairwa G, Jung WH, Kronstad JW. 2017. Iron acquisition in fungal pathogens of humans. Metallomics 9:215–227.

9. Vartivarian SE, Anaissie EJ, Cowart RE, Sprigg HA, Tingler MJ, Jacobson ES. 1993. Regulation of cryptococcal capsular polysaccharide by iron. J Infect Dis. 167:186–190.

10. Jung WH, Sham A, Lian T, Singh A, Kosman DJ, Kronstad JW. 2008. Iron source preference and regulation of iron uptake in *Cryptococcus neoformans*. PLoS Pathog 4:e45.

11. Hood MI, Skaar EP. 2012. Nutritional immunity: transition metals at the pathogen-host interface. Nat Rev Microbiol 10:525–537.

12. Jung WH, Hu G, Kuo W, Kronstad JW. 2009. Role of ferroxidases in iron uptake and virulence of *Cryptococcus neoformans*. Eukaryot Cell 8:1511–1520.

13. Cadieux B, Lian T, Hu G, Wang J, Biondo C, Teti G, Liu V, Murphy ME, Creagh AL, Kronstad JW. 2013. The mannoprotein Cig1 supports iron acquisition from heme and virulence in the pathogenic fungus *Cryptococcus neoformans*. J Infect Dis 207:1339–1347.

14. Saikia S, Oliveira D, Hu G, Kronstad J. 2014. Role of ferric reductases in iron acquisition and virulence in the fungal pathogen *Cryptococcus neoformans*. Infect Immun 82:839–850.

15. Jung WH, Sham A, White R, Kronstad JW. 2006. Iron regulation of the major virulence factors in the AIDS-associated pathogen *Cryptococcus neoformans*. PLoS Biol 4:e410.

16. Jung, W. H., Saikia, S., Hu, G., Wang, J., Fung, C. K., D’Souza, C., et al. (2010) HapX positively and negatively regulates the transcriptional response to iron deprivation in *Cryptococcus neoformans*. PLoS Pathog 6: e1001209.

17. Schrettl M, Kim HS, Eisendle M, Kragl C, Nierman WC, Heinekamp T, Werner ER, Jacobsen I, Illmer P, Yi H, Brakhage AA, Haas H. 2008. SreA-mediated iron regulation in *Aspergillusfumigatus*. Mol Microbiol 70:27–43.

18. Harrison KA, Marzluf GA. 2002. Characterization of DNA binding and the cysteine rich region of SRE, a GATA factor in Neurospora crassa involved in siderophore synthesis Biochemistry. 41:15288–15295.

19. Pelletier B, Beaudoin J, Mukai Y, Labbé S. 2002. Fep1, an iron sensor regulating iron transporter gene expression in *Schizosaccharomyces pombe*. J Biol Chem 277:22950–22958.

20. Haas H, Zadra I, Stöffler G, Angermayr K. 1999. The *Aspergillus nidulans* GATA factor SREA is involved in regulation of siderophore biosynthesis and control of iron uptake. J Biol Chem 274:4613–4619.

21. An Z, Mei B, Yuan WM, Leong SA. 1997. The distal GATA sequences of the sid1 promoter of *Ustilago maydis* mediate iron repression of siderophore production and interact directly with Urbs1, a GATA family transcription factor. EMBO J. 16:1742–1750.

22. Yamaguchi-Iwai Y, Dancis A, Klausner RD. 1995. AFT1: a mediator of iron regulated transcriptional control in Saccharomyces cerevisiae. EMBO J 14:1231–1239.

23. Blaiseau PL, Lesuisse E, Camadro JM. 2001. Aft2p, a novel iron-regulated transcription activator that modulates, with Aft1p, intracellular iron use and resistance to oxidative stress in yeast. J Biol Chem 276:34221–34226.

24. Li H, Outten CE. 2012. Monothiol CGFS glutaredoxins and BolA-like proteins: [2Fe-2S] binding partners in iron homeostasis. Biochemistry 51:4377–4389.

25. Outten CE, Albetel AN. 2013. Iron sensing and regulation in *Saccharomyces cerevisiae:* Ironing out the mechanistic details. Curr Opin Microbiol 16:662–668.

26. Labbé S, Khan MG, Jacques JF. 2013. Iron uptake and regulation in *Schizosaccharomyces pombe*. Curr Opin Microbiol 16:669–676.

27. Encinar del Dedo J, Gabrielli N, Carmona M, Ayté J, Hidalgo E. 2015. A Cascade of Iron-Containing Proteins Governs the Genetic Iron Starvation Response to Promote Iron Uptake and Inhibit Iron Storage in Fission Yeast. PLoS Genet 11: e1005106.

28. Ojeda L, Keller G, Muhlenhoff U, Rutherford JC, Lill R, Winge DR. 2006. Role of glutaredoxin-3 and glutaredoxin-4 in the iron regulation of the Aft1 transcriptional activator in *Saccharomyces cerevisiae*. J Biol Chem 281:17661–17669.

29. Jbel M, Mercier A, Labbé S. 2011. Grx4 monothiol glutaredoxin is required for iron limitation-dependent inhibition of Fep1. Eukaryot Cell 10:629–645.

30. Kim KD, Kim HJ, Lee KC, Roe JH. 2011. Multi-domain CGFS-type glutaredoxin Grx4 regulates iron homeostasis via direct interaction with a repressor Fep1 in fission yeast. Biochem Biophys Res Commun 408:609–614.

31. Ueta R, Fujiwara N, Iwai K, Yamaguchi-Iwai Y. 2012. Iron-induced dissociation of the Aft1p transcriptional regulator from target gene promoters is an initial event in iron-dependent gene suppression. Mol Cell Biol 32:4998–5008.

33. Vachon P, Mercier A, Jbel M, Labbé S. 2012. The monothiol glutaredoxin Grx4 exerts an iron-dependent inhibitory effect on Php4 function. Eukaryot Cell 11:806–819.

34. Dlouhy AC, Beaudoin J, Labbé S, Outten CE. 2017. *Schizosaccharomyces pombe* Grx4 regulates the transcriptional repressor Php4 via [2Fe-2S] cluster binding. Metallomics 9:1096–1105.

35. Meyer Y, Buchanan BB, Vignols F, Reichheld JP. 2009. Thioredoxins and glutaredoxins: unifying elements in redox biology. Annu Rev Genet 43:335–367.

36. Rouhier N, Couturier J, Johnson MK, Jacquot JP. 2010. Glutaredoxins: roles in iron homeostasis. Trends Biochem Sci 35:43–52.

37. Hu G, Caza M, Cadieux B, Bakkeren E, Do E, Jung WH, Kronstad JW. 2015. The endosomal sorting complex required for transport machinery influences haem uptake and capsule elaboration in *Cryptococcus neoformans*. Mol Microbiol 96:973–992.

38. Mercier A, Watt S, Bahler J, Labbé S. 2008. Key function for the CCAAT-binding factor Php4 to regulate gene expression in response to iron deficiency in fission yeast. Eukaryot Cell 7:493–508.

39. Mercier A, Labbé S. 2009. Both Php4 function and subcellular localization are regulated by iron via a multistep mechanism involving the glutaredoxin Grx4 and the exportin Crm1. J Biol Chem 284:20249–20262.

40. Szklarczyk D, Morris JH, Cook H, Kuhn M, Wyder S, Simonovic M, Santos A, Doncheva NT, Roth A, Bork P, Jensen LJ, von Mering C. 2017. The STRING database in 2017: quality-controlled protein-protein association networks, made broadly accessible. Nucleic Acids Res. 45:D362–368.

41. Zhang C. 2014. Essential functions of iron-requiring proteins in DNA replication, repair and cell cycle control. Protein Cell. 5:750–760.

42. Puig S, Ramos-Alonso L, Romero AM, Martínez-Pastor MT. 2017. The elemental role of iron in DNA synthesis and repair. Metallomics 9:1483–1500.

43. Muhlenhoff U, Molik S, Godoy JR, Uzarska MA, Richter N, Seubert A, Zhang Y, Stubbe J, Pierrel F, Herrero E, Lillig CH, Lill R. 2010. Cytosolic monothiol glutaredoxins function in intracellular iron sensing and trafficking via their bound iron-sulfur cluster. Cell Metab 12:373–385.

44. Ueta R, Fujiwara N, Iwai K, Yamaguchi-Iwai Y. 2007. Mechanism underlying the iron-dependent nuclear export of the iron-responsive transcription factor Aft1p in *Saccharomyces cerevisiae*. Mol Biol Cell. 18:2980–2990.

45. McNabb DS, Xing Y, Guarente L. 1995. Cloning of yeast HAP5: a novel subunit of a heterotrimeric complex required for CCAAT binding. Genes Dev. 9:47–58.

46. Schrettl M, Beckmann N, Varga J, Heinekamp T, Jacobsen ID, Jochl C, Moussa TA, Wang S, Gsaller F, Blatzer M Werner ER, Niermann WC, Brakhage AA, Haas H. 2010. HapX-mediated adaption to iron starvation is crucial for virulence of *Aspergillus fumigatus*. PLoS Pathog 6:e1001124.

47. Gsaller F, Hortschansky P, Beattie SR, Klammer V, Tuppatsch K, Lechner BE, Rietzschel N, Werner ER, Vogan AA, Chung D, Mühlenhoff U, Kato M, Cramer RA, Brakhage AA, Haas H. 2014. The Janus transcription factor HapX controls fungal adaptation to both iron starvation and iron excess. EMBO J. 33:2261–2276.

48. Hsu PC, Yang CY, Lan CY 2011. *Candida albicans* Hap43 is a repressor induced under low-iron conditions and is essential for iron-responsive transcriptional regulation and virulence. Eukaryot Cell 10:207–225.

49. Pham K, Pal R, Qu Y, Liu X, Yu H, Shiao SL, Wang X, O’Brian Smith E, Cui X, Rodney GG, Cheng N. 2015. Nuclear glutaredoxin 3 is critical for protection against oxidative stress-induced cell death. Free Radic Biol Med. 85:197–206.

50. Pujol-Carrion N, de la Torre-Ruiz MA. 2010. Glutaredoxins Grx4 and Grx3 of *Saccharomyces cerevisiae* play a role in actin dynamics through their Trx domains, which contributes to oxidative stress resistance. Appl Environ Microbiol. 76:7826–7835.

51. Chaves GM, da Silva WP. 2012. Superoxide dismutases and glutaredoxins have a distinct role in the response of *Candida albicans* to oxidative stress generated by the chemical compounds menadione and diamide. Mem Inst Oswaldo Cruz. 107:998–1005.

52. Zhang LB, Tang L, Ying SH, Feng MG. 2016. Regulative roles of glutathione reductase and four glutaredoxins in glutathione redox, antioxidant activity, and iron homeostasis of *Beauveria bassiana*. Appl Microbiol Biotechnol 100:5907–5917.

53. Zhang D, Dong Y, Yu Q, Kai Z, Zhang M, Jia C, Xiao C, Zhang B, Zhang B, Li M. 2017. Function of glutaredoxin 3 (Grx3) in oxidative stress response caused by iron homeostasis disorder in *Candida albicans*. Future Microbiol 12:1397–1412.

54. Toffaletti DL, Rude TH, Johnston SA, Durack DT, Perfect JR. 1993. Gene transfer in *Cryptococcus neoformans* by use of biolistic delivery of DNA. J Bacteriol. 175:1405–1411.

55. Davidson RC, Blankenship JR, Kraus PR, de Jesus Berrios M, Hull CM, D’Souza C, Wang P, Heitman J. 2002. A PCR-based strategy to generate integrative targeting alleles with large regions of homology. Microbiology 148:2607–2615.

56. Yu JH, Hamari Z, Han KH, Seo JA, Reyes-Domínguez Y, Scazzocchio C. 1994. Doublejoint PCR: a PCR-based molecular tool for gene manipulations in filamentous fungi. Fungal Genet Biol 41:973–981.

57. Bolger, A.M., Lohse, M. and Usadel, B. 2014. Trimmomatic: a flexible trimmer for Illumina sequence data. Bioinformatics 30:2114–2120.

58. Kim, D., Langmead, B. and Salzberg, S.L. 2015. HISAT: a fast spliced aligner with low memory requirements. Nat Methods 12:357–360.

59. Li H, Handsaker B, Wysoker A, Fennell T, Ruan J, Homer N, Marth G, Abecasis G, Durbin R; 1000 Genome Project Data Processing Subgroup. 2009. The Sequence Alignment/Map format and SAMtools. Bioinformatics 25:2078–2079.

60. Anders S, Pyl PT, Huber W. 2015. HTSeq–a Python framework to work with high-throughput sequencing data. Bioinformatics 31:166–169.

61. Robinson MD, McCarthy DJ, Smyth GK. 2010. edgeR: a Bioconductor package for differential expression analysis of digital gene expression data. Bioinformatics 26:139–140.

62. Trapnell C, Roberts A, Goff L, Pertea G, Kim D, Kelley DR, Pimentel H, Salzberg SL, Rinn JL, Pachter L. 2012. Differential gene and transcript expression analysis of RNA-seq experiments with TopHat and Cufflinks. Nature Protocols 7:562–578.

63. Jones P, Binns D, Chang HY, Fraser M, Li W, McAnulla C, McWilliam H, Maslen J, Mitchell A, Nuka G, Pesseat S, Quinn AF, Sangrador-Vegas A, Scheremetjew M, Yong SY, Lopez R, Hunter S. 2014. InterProScan 5: genome-scale protein function classification. Bioinformatics 30: 1236–1240.

64. Falcon S, Gentleman R. 2007 Using GOstats to test gene lists for GO term association. Bioinformatics 23:257–258.

65. Wickham, H. 2009. ggplot2: elegant graphics for data analysis, Springer-Verlag, New York, NY.

